# Uncovering Heterogeneous Effects via Localized Feature Selection

**DOI:** 10.1101/2025.06.03.657761

**Authors:** Xiaoxia Liu, Jiaqi Gu, Zhaomeng Chen, Benjamin Chu, Linxi Liu, Tim Morrison, Robert R. Butler, Jacob Edelson, Jinzhou Li, Frank M. Longo, Hua Tang, Iuliana Ionita-laza, Chiara Sabatti, Emmanuel Candès, Zihuai He

## Abstract

Identifying features that interact to trigger disease, while accounting for heterogeneity across diverse populations, is essential for the development of precision and targeted medicine. Despite the availability of vast and complex health-related datasets, most existing works focus on identifying disease-associated features at the population level or within a few subpopulations, often overlooking individual-level heterogeneity within these groups. To address this limitation, we propose a novel framework that utilizes localized test statistics to identify disease-associated features tailored to individual profiles. Our method leverages the recently developed knockoffs methodology to control the noise level of the selection set so that the results are replicable. Moreover, it allows for the discovery of hidden heterogeneous effects within the data, as demonstrated in an application to single-cell RNA sequencing data for Alzheimer’s disease. By aggregating localized feature selection results, our framework also enables powerful population-level feature selection. Our framework provides a powerful tool for exploratory studies of precision medicine, offering the potential to generate novel hypotheses for confirmatory biological experiments.

## 1. Introduction

Tailoring treatments to patients based on their individual characteristics is a key goal in medical practice. To achieve this goal, one main theme in precision medicine is identifying features that interact to trigger a disease cascade while taking patient heterogeneity into account. The discoveries can provide mechanistic insights and potentially lead to the development of personalized therapy that delivers the optimal treatment to the right patient at the right time.

To date, most studies that tackle heterogeneity focus on the identification of disease-associated features at the level of discrete subpopulations, such as those defined by age, sex, and population ancestry ^1–5^. However, ignoring heterogeneity within each category can induce significant bias. For example, using self-reported ancestry as a genetic category overlooks the continuous nature of genetic ancestries, which may lead to misleading conclusions, particularly for individuals with uncommon admixed ancestry proportions ^6–9^. The recent discovery that polygenic score accuracy varies across the continuum of genetic ancestry also highlights the need to shift from discrete genetic ancestry clusters to a continuum of genetic ancestries ^10^.

Compared to feature selection at the population level, most applications can benefit more from “localized feature selection”. In medical contexts, localized feature selection enables the discovery of disease-associated features tailored to individual profiles or to subgroups derived from high-dimensional data. In the machine learning literature, various localized feature importance scores have been developed to quantify the effect of a feature on the disease outcome in a sophisticated model, referred to as explainable artificial intelligence (XAI), including but not limited to gradient backpropagation methods (e.g. gradients, integrated gradients, expected gradients) and “explaining by removing” methods (e.g. SAGE, SHAP, LIME). While those importance scores do characterize relative localized importance of features, they do not provide a feature selection set with error rate control, which is required for reproducible science.

The framework we introduce in this paper enables localized selection of features whose “importance” may vary across different samples. By leveraging nonlinear models, we construct localized test statistics that utilize information from the entire study sample while emphasizing relationships between explanatory variables and the outcome within a neighborhood of the covariate space. We leverage the recently developed knockoffs methodology to control the noise level of the selection set, improving the replicability of results. Examining how selection sets derived from these test statistics vary across the study sample can uncover hidden population substructures. Moreover, by computing these localized test statistics on new feature values, we can assess feature importance specific to new observations.

Our framework is based on model-X knockoffs ^11^, a procedure that performs feature selection with rigorous error rate control in high-dimensional settings. Unlike classical approaches that select important features through stagewise model fitting ^12–15^ or regularization ^16–20^, model-X knockoffs generates a set of features, called knockoffs, which serve as negative controls. These knockoffs are designed to be statistically indistinguishable from the corresponding original features and are constructed without reference to the response. This, combined with almost any feature importance scores and the knockoffs filter ^11^, allows the analyst to identify important variables with FDR control. Like model-X knockoffs, our framework is also compatible with a large class of machine learning algorithms, including generalized linear models, random forests, and neural networks with symmetric structures. It can also flexibly integrate recent advances in explainable artificial intelligence (XAI), which provide novel localized feature importance scores to understand the features that drive the predictions. By aggregating the localized scores derived from knockoffs, our method allows for powerful selection of otherwise unrecognized population features. We also develop a derandomization strategy that aggregates selection sets obtained from multiple runs to further improve replicability of the feature selection. The advantages of our framework are demonstrated through various simulations and an application to single-cell RNA sequencing data for Alzheimer’s disease, where applying our procedure manages to identify cell subtypes with different gene importance patterns, which cannot be distinguished by unsupervised embeddings using the feature matrix. This is of note since differential properties and behaviors across cellular subtypes within cell populations are receiving increasing recognition within the context of AD pathology progression ^21^.

## 2. Results

### A. Overview of the proposed localized feature selection

Consider a dataset 𝔻 = {𝕩 = [***x***_1_, …, ***x***_*n*_]^*T*^, ***Y*** = (*y*_1_, …, *y*_*n*_)^*T*^} that contains *n* individuals, where 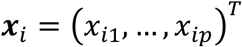 and *y* represent the *p* features and the response of the *i*-th individual, respectively (*i* = 1, … *n*). Our goal is to select key features related to the outcome, recognizing that their importance may differ across samples. Specifically, for a given vector of feature values *x*, we aim to find a subset of features *ℛ*_***x***_ ⊂ {1, …, *p*} that are associated with the outcome while making sure that the error rate is less than a predefined level.

Although various XAI methods have been developed to assess feature contributions (e.g. gradients, SAGE, SHAP, LIME), they often struggle with valid statistical inference and error rate control, hindering replicability. To address this, we adapt model-X knockoffs (11), a method that can filter out unwanted noise in the feature importance and rigorously controls the FDR of the selection set *ℛ*_***x***_. **Figure 1** summarizes the workflow of our proposed method.

**Figure 1:**
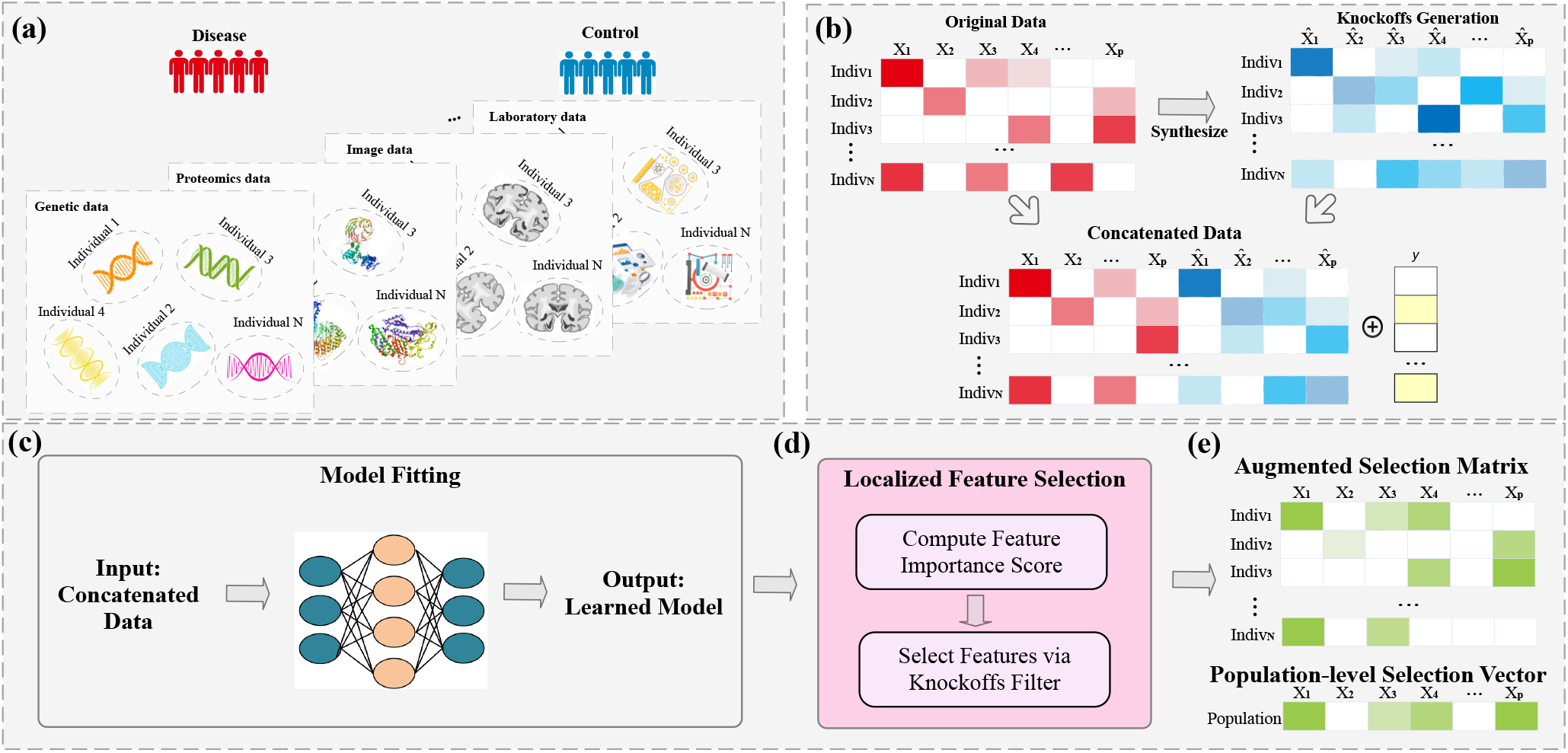
Graphical illustration of the proposed localized feature selection workflow.

For each feature, we construct a set of knockoff data consisting of a set of synthetic controls (**Figure 1 (b)**). Both the original and knockoff data are input into a machine learning method (**Figure 1 (c)**). Knockoffs serve as control features during training and subsequently help identify variables that truly explain the response variable.

Once the model is fitted, we use feature importance scores to measure each feature’s localized contribution to the response compared to its knockoff copy. A knockoff filter selects features with controlled errors at different target FDR threshold values (e.g. 10%, 20%; **Figure 1 (d)**). The results across all data points A are summarized as an augmented selection matrix (**Figure 1 (e)**). This matrix can be used in downstream exploratory analyses to uncover hidden data structures. Our framework also supports powerful population feature selection by aggregating localized inference results.

We describe the details, including mathematical formulation, construction of knockoffs, model fitting, localized feature importance calculation, feature selection, and derandomization (knockoffs are randomly generated) in the Method Section (Section 3).

### B. Localized feature selection reveals heterogeneous effects

#### Description of the dataset

We apply our framework to a single-cell RNA sequencing data (scRNA-seq) collected and preprocessed by Stanford/VA/NIA Aging Clinical Research Center (ACRC) ^22^. This dataset includes single-nucleus transcriptomes from 143,793 cells derived from the hippocampal regions of 17 individuals (8 controls and 9 AD cases) and the cortical regions of 8 individuals (4 controls and 4 AD cases, a subset of the 17 above). For each cell, the scRNA-seq data records transcriptome counts for 23,537 genes. Given the critical role of the microglia cell type in neuroinflammation and its potential involvement in AD pathology ^23, 24^, we focus on 3,373 microglia, with 2,526 derived from the hippocampus and 847 from the cortex. We then remove genes that are rarely expressed in microglia by filtering out all genes with no more than 5% nonzero counts, resulting in a transcriptome count data A for *p* = 2,955 genes across *n* = 3,373 cells. For the *i*-th cell, we define the response *y*_*i*_ as 1 if the cell comes from an AD case and 0 otherwise, resulting in 2,231 AD cells and 1,142 control cells. The goal of this analysis is to characterize heterogeneous effect of genes on AD at individual cell level and uncover hidden patterns in the data. The number of microglia from each individual and brain region is provided in **Supplementary Table 1**.

#### Implementation of the method

We first construct knockoffs 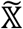 and apply stringent quality control to ensure that the knockoffs obey the exchangeability property, i.e. they are statistically indistinguishable from the original features (see Section S1.A for details).^11,25^ After generating knockoffs 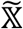, we construct localized test statistics by fitting a quadratic regression model *h*(⋅) between ***Y*** and the concatenated data matrix 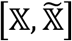 as detailed in Section 3.C and Section S1.B of the supplementary material.

To enhance the stability and replicability of the analysis, we implement the derandomization strategy described in Section 3.E. Specifically, we perform *M* = 50 runs of our procedure with a target FDR level *α* = 0.2, using the quadratic model (S1) with transcriptome counts of the 2,955 genes, batch indices (a categorical feature with 4 levels), and the original region (a categorical feature with 2 levels) of the cells as features. To assess the benefit of employing a quadratic model over a linear model, we compare the average cross-validation *R*^2^ of the fitted quadratic model (0.4011) with the average cross-validation *R*^2^ of the fitted lasso linear model (0.3260) over *M* = 50 runs. The quadratic model explains 23% more variance in the response *y*_*i*_’s, suggesting the presence of nonlinear relations between gene expressions and AD. For each cell *i* = 1, …, 3373, we compute the derandomized selection set 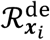 and the population-level selection set 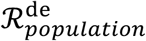 according to the procedures described in Sections 3.C to 3.E. We generate the augmented selection matrix ***V***_𝔻_(as defined in Section 3.E) across all microglia. The genes selected in 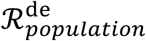 and their inclusion rates are presented in Supplementary Table 2. To quantify the stability of the sets 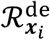 and 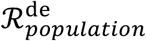, we reapply our procedure with the derandomization strategy for another 30 runs and compute the average Jaccard similarity of 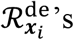 and 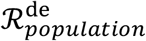 across these runs. As shown in Supplementary Figure 6, the average Jaccard similarities for most 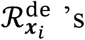 exceeds 0.9, while the average Jaccard similarity of 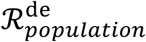 reaches 0.909, demonstrating the stability of our method in analyzing the scRNA-seq data.

### UMAP analysis of augmented selection matrix vs. existing alternatives

We further analyze the augmented selection matrix ***V***_𝔻_, computed as described in Section 3.E, to uncover potential subgroup structures. Specifically, we perform the UMAP (Uniform Manifold Approximation and Projection ^39^) transformation on ***V***_𝔻_ and apply *k*-means clustering (*k* = 6) on the UMAP embeddings of the 3,373 cells. The UMAP embeddings are displayed in Figure 2 (c). The results reveal a discrete clustering pattern among microglia with at least six distinct groups. We further annotated the cells by their brain regions and found that, without pre-specifying subpopulations, the method discovered that groups 1 and 2 consist exclusively of cortex cells, while groups 3–6 consist exclusively of hippocampus cells. Notably, this separation may reflect differential AD progression established between the two brain regions ^21, 26, 27^ and is not apparent in the UMAP embeddings of the 3,373 cells using either the original data matrix A (Figure 2 (a)) or the knockoff feature importance scores 𝕏 (Figure 2 (b)) as input. While previous studies have identified regional neuronal populations purely via transcriptomic data^28–32^, this is a novel separation of microglia without pre-hoc information about brain region. This demonstrates that our localized feature selection approach can uncover a distinct and previously hidden clustering structure among microglia.

**Figure 2.**
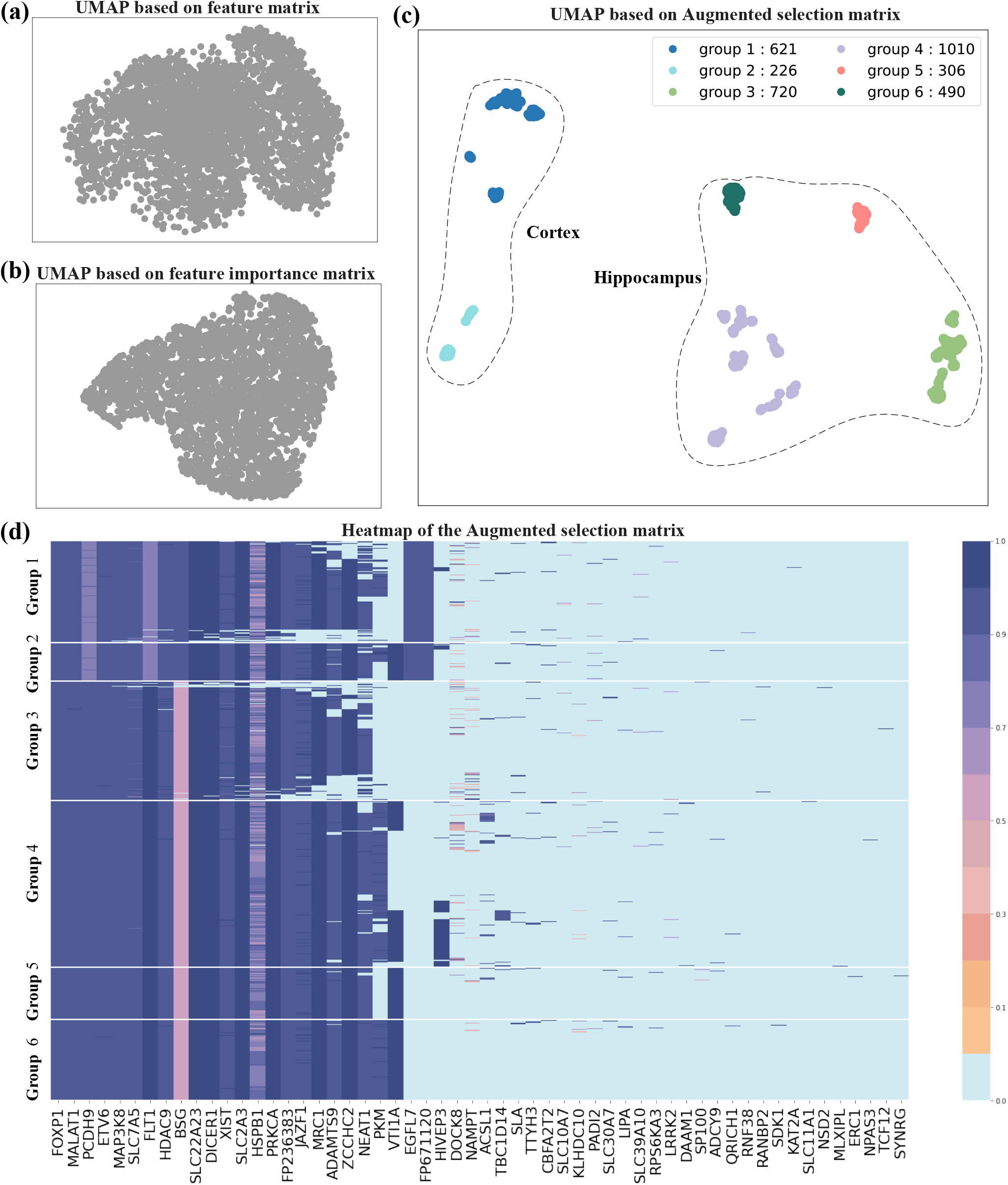
(a) UMAP embeddings generated from the data matrix 𝕏. (b) UMAP embeddings generated from the matrix 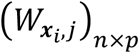 of localized feature importance scores. (c) UMAP embeddings generated from the augmented selection matrix ***V***_𝔻_. (d) Heatmap of the augmented selection matrix ***V***_𝔻_, where a light blue grid at the *i*-th row and *j*-th column indicates that 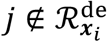.

### Exploring analysis results to generate novel scientific hypotheses

Based on the six identified groups, we rearrange the indices of cells according to their group labels and visualize the augmented selection matrix *V*_𝔻_ in Figure 2 (d). For better exposition, we present the 56 genes that are included in at least one localized selection set 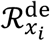. It is clear that the localized selection sets 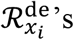 differ for cells in different groups. Comparing groups 1 and 2 (from the cortex region) with groups 3, 4, 5 and 6 (from the hippocampus region), we observe distinct gene effects for genes such as PCDH9, FLT1, BSG, EGFL7, and transcript FP671120. Interestingly, PCDH9 is reflective of a higher AD feature importance in hippocampal cells than in cortical cells, which agrees with Chen et al ^33^ who found a PCDH9^high^ microglial subtype that predominantly localized to the hippocampus, having isolated both brain regions separately. Further Chen et al tested the differential expression of these PCDH9^high^ microglia in cortex versus hippocampus and found upregulation of BSG expression in the cortex echoing its higher feature importance in our group 1 & 2 cortical data. Additionally, Young et al ^34^ had previously identified a FLT1 expressing microglial subtype, associated with cell proliferation and chemotaxis, though they lacked a hippocampal sample for direct comparison.

Even among cells within the same brain region, our method identifies heterogeneous gene effect patterns that are unrecognized by other approaches. For example, although both groups 1 and 2 are from the cortex, our method reveals that the AD feature importance of VTI1A strongly discriminates group 2 cells. VTI1A plays an important role in glial and synaptic vesicular signaling ^35^, and GWAS loci for VTI1A have been associated with increased glioma risk ^36^, suggesting possible functional role for this group 2 cortical microglia subpopulation that is AD-relevant. Distinctions among subpopulations 3, 4, 5 and 6 are also evident based on the differential AD feature importance of PKM, a known marker of microglia activation ^37, 38^. These findings illustrate that our localized feature selection method effectively identifies biologically relevant gene effects in plausible subpopulations of microglia that can be used to further stratify and investigate AD-relevant changes not apparent via conventional transcriptomic analysis without pre-hoc knowledge of regional or functional groupings.

### C. Simulation studies

To evaluate the performance of the proposed localized feature selection framework, we conduct extensive simulations to demonstrate:

- The FDR control of both the localized selection and the population-level selection.
- The ability of the localized selection sets to capture heterogeneous effects.
- The power enhancement of the population-level selection by aggregating localized test statistics compared to feature selection based on a linear model.
- The improved replicability by derandomizing the selection set.

We simulate features ***G*** = (*G*_1_, …, *G*_100_)^*T*^ of each individual from the multivariate normal distribution

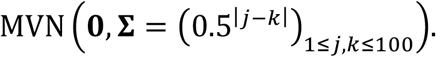

To emulate the inclusion of demographic covariates that contribute to the heterogeneity of real genomic data, we further simulate three independent features

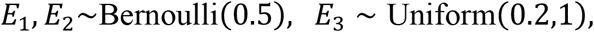

to mimic two ancestry categories (European American, EUR, if *E*_1_ = 0 and African American, AFA, if *E*_1_=1), two sex categories (male if *E*_2_ = 0 and female if *E*_2_=1), and age (equals *E*_3_ × 100). Throughout all simulation experiments, we generate the response *Y* from

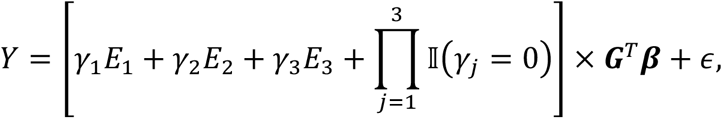

where ϵ ~*N*(0,1) is independent of everything else.

We consider four scenarios to investigate the impact of different effect sizes attributed to the covariates (*E*_1_, *E*_2_, *E*_3_) on the performance of localized feature selection:

- Scenario 1: 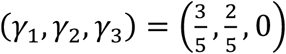;
- Scenario 2: (*γ*_1_, *γ*_2_, *γ*_3_) = (0,0,1);
- Scenario 3: 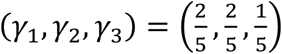;
- Scenario 4: (*γ*_1_, *γ*_2_, *γ*_3_) = (0,0,0).

By doing so, we simulate scenarios in which features have a stronger impact on the response in certain groups compared to others. For example, scenario 1 highlights the impact of ancestry and sex on the heterogeneity of genetic variants’ contributions to the variation of *Y*, while scenario 2 focuses on the age-dependent genetic effect. Throughout all four scenarios, we let 20 randomly selected entries of β be 0.38 or −0.38 with equal probability and set the remaining entries to 0. For each scenario, keeping the parameter values fixed, we simulate independently 100 replicates of the dataset 𝔻 = {𝕏, ***Y***}, each with a sample size of 1000. We then perform localized feature selection over the *p* = 103 features on these datasets. For a comprehensive investigation of how localized selection set *ℛ*_***x***_ captures effect heterogeneity across different neighborhoods, we compute *ℛ*_***x***_’s for 2000 evaluation points (***x***) in the sample space across all 100 replicates.

### D. Simulation results: localized feature selection

We first investigate the performance of our method in controlling FDR and identifying important features under different scenarios. After generating knockoffs 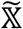, we construct localized test statistics by fitting a quadratic regression model *h*(⋅) between ***Y*** and the concatenated data matrix 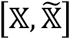 as detailed in Section S1.B of the supplementary material. Specifically, for each localized selection set *ℛ*_***x***_ corresponding to an evaluation point ***x***, we report the average across 100 replicates of the following quantities:

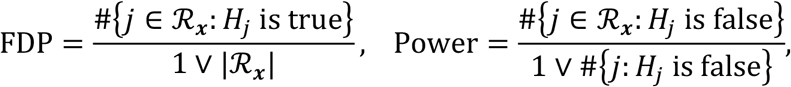

where *H*_*j*_ ‘s are the conditional independence hypotheses:

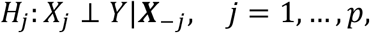

where ***X*** =(*X*_1_, …, *X*_*j*–1_, *X*_*j*+1_, …, *X*_*p*_)^*T*^ records all features except *X*. A non-null means that the *j*-th feature has predictive power beyond that afforded by all the other features. *ℛ*_***x***_ is derived with a target FDR level of *α* = 0.1. Figure 3 presents the empirical FDR and power of the localized selection sets *ℛ*_***x***_’s under different scenarios, where each point corresponds to *ℛ*_***x***_ for one of the 2,000 evaluation points. Under all 4 scenarios, the empirical FDR of all the *ℛ*_***x***_’s are under the target level *α* = 0.1, which validates the FDR control of localized selection sets. Moreover, there are varying patterns in the power of *ℛ*_***x***_’s across different scenarios. As shown in Figure 3 (a), under scenario 1, the power of *ℛ*_***x***_’s varies among different ancestry and gender profiles (*E*_1_, *E*_2_) while remaining consistent with respect to age. This is because the impact of an important feature *G*_*j*_ on the response *Y* is most significant for AFA women, indicating strong localized evidence against null hypotheses. In contrast, for EUR men (with *E*_1_ = *E*_2_ = 0), the variation of *G*_*j*_ does not affect the response *Y*, resulting in no localized evidence against false hypotheses and, consequently, neglectable power. Under scenario 2, where only age (*E*_3_) determines the strength of localized evidence against *H*_*j*_’s, Figure 3 (b) shows that the power of *ℛ*_***x***_’s varies across individuals of different ages, while no significant power difference exists among different profiles of ancestry and gender profiles (*E*_1_, *E*_2_). Unlike EUR men (with *E*_1_ = *E*_2_ = 0), who had no localized evidence against *H*_*j*_’s under scenario 1, weak localized evidence against *H*_*j*_’s still exists in neighborhoods of ***x*** where *E*_3_ *is* close to 0.2, resulting in non-zero power for the corresponding *ℛ*_***x***_. Figure 3 (c) presents the power of *ℛ*_***x***_’s under scenario 3, where all three covariates impact the strength of localized evidence against false *H*_*j*_’s, with larger values of *E*_1_, *E*_2_, *E*_3_ corresponding to stronger localized evidence. As with scenarios 1 and 2, the power of *ℛ*_***x***_’s is higher for elder AFA women (with *E*_1_ = *E*_2_ = 1 and a large *E*_3_). In scenario 4, shown in Figure 3 (d), where there is no heterogeneity in localized evidence against false *H*_*j*_’s among different individuals, the power of *ℛ*_***x***_’s does not differ significantly among different ***x***’s.

**Figure 3.**
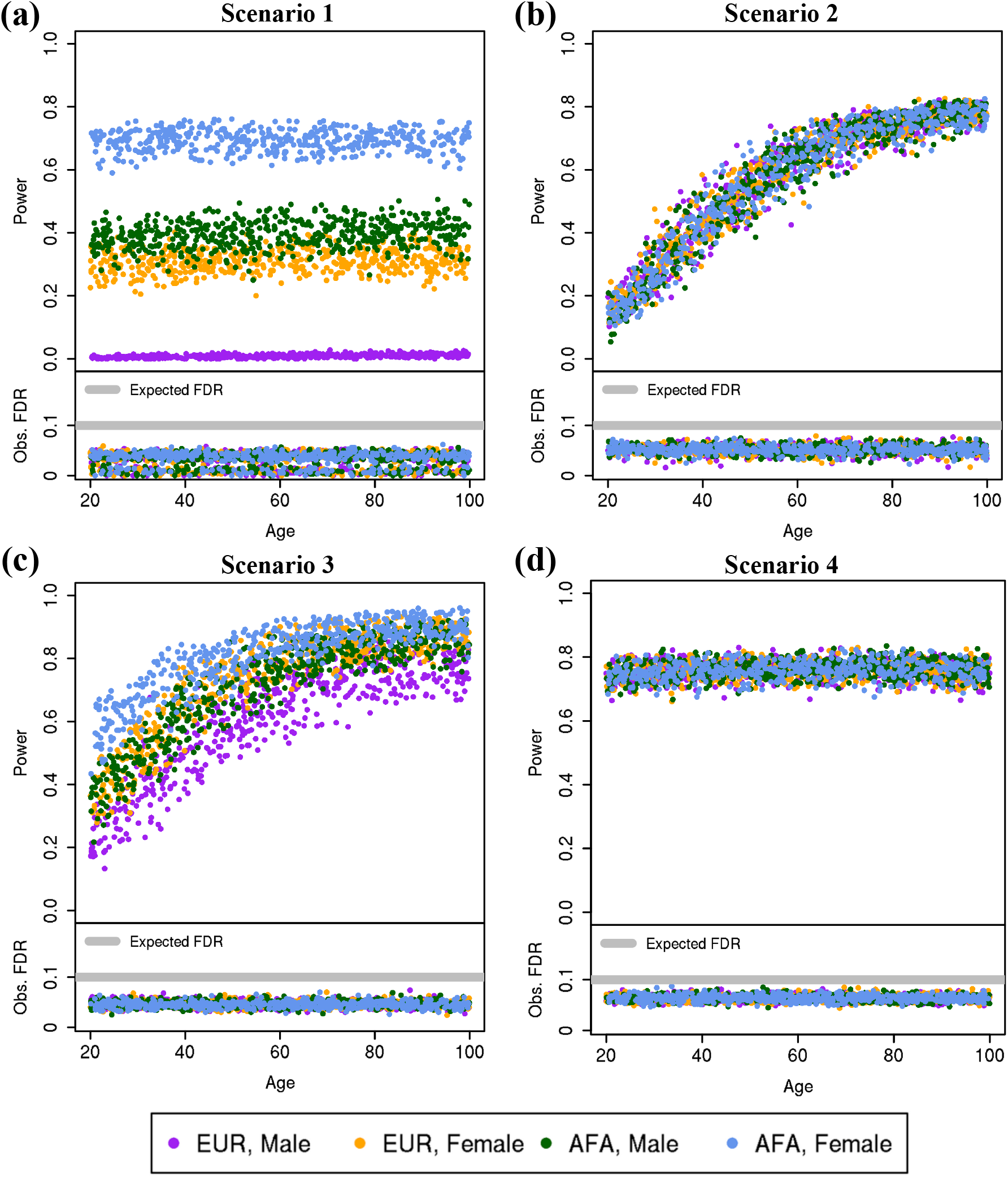
Average (across 100 replicates) power and FDP of the localized selection sets *ℛ*_***x***_’s (with respect to the target FDR level *α* = 0.1) for each of the 2000 evaluation points across the four scenarios. Each dot corresponds to one *ℛ*_***x***_ and it is colored according to the value of the ancestry and sex feature in ***x***.

The patterns observed under different scenarios are also reflected in the UMAP visualizations shown in Figure 4. Specifically, we use the augmented selection matrix, obtained by concatenating the augmented selection vectors ***v***_***x***_’s of the 2000 evaluation points from one replicate, as the input to the UMAP algorithm. Points are colored according to the values of age, ancestry, and sex feature, with different color mappings used in the two displays. In Figure 4 (a), the ***v***_***x***_’s of different ancestry and gender profiles are clearly separated in the projections, while in Figure 4 (b), the projections align according to age. Scenario 3 (Figure 4 (c)) demonstrates separation based on ancestry, gender, and age, whereas in scenario 4 (Figure 4 (d)), projections of all ***v***_***x***_’s are well-mixed, consistent with the results in Figure 3. This distinct alignment of UMAP projections suggests that our procedure effectively captures the driving factors of effect heterogeneity.

**Figure 4.**
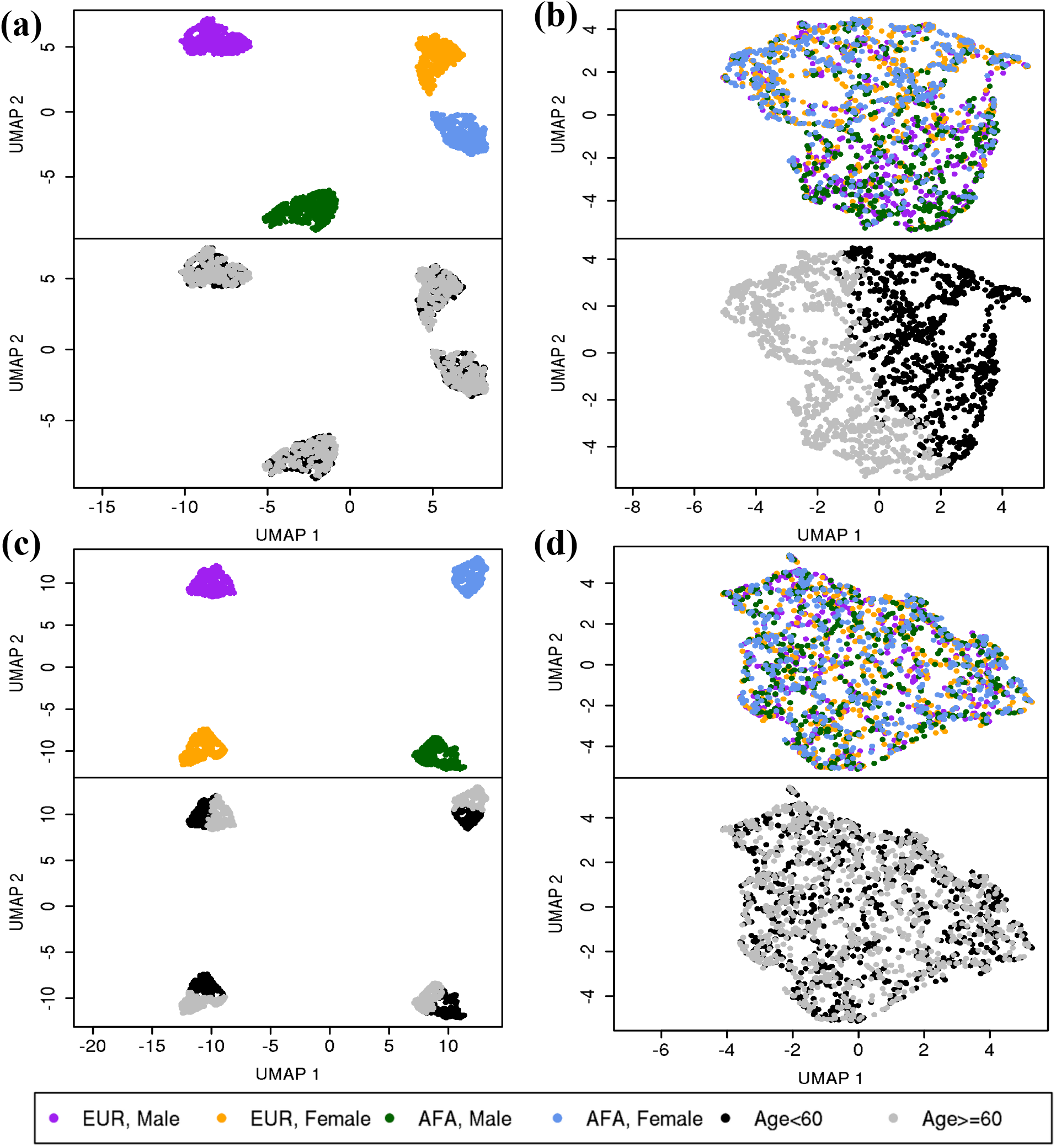
UMAP visualization of the augmented selection vectors ***v***_***x***_’s from one replicate under different scenarios. The upper panel of each subfigure is colored according to ancestry and gender profiles (*E*_1_, *E*_2_), while the lower panel is colored according to whether the age exceeds 60 (the mean and median of age).

### E. Simulation results: population-level feature selection

In addition to generating the localized feature selection sets *ℛ*_***x***_’s, our framework also facilitates powerful population-level feature selection by aggregating the localized inference results, as introduced in Section 3.D. Specifically, by summing up *W*_***x***,*j*_’s from different evaluation points ***x***’s, we effectively summarize evidence against the nulls while accounting for heterogeneous patterns and varying strengths. To illustrate this, we visualize the empirical power and FDR associated with:

1. *ℛ*_population_, obtained by aggregating *W*_***x***,*j*_’s across all evaluation points;
2. *ℛ*_young_eur_men_, obtained by aggregating *W*_***x***,*j*_’s across all evaluation points corresponding to EUR men under 60 (*E*_1_ = *E*_2_ = 0, *E*_3_ *<* 0.6);
3. *ℛ*_old_afa_women_, obtained by aggregating *W*_***x***,*j*_’s across all evaluation points corresponding to AFA women aged above 60 (*E*_1_ = *E*_2_ = 1, *E*_3_ *>* 0.6);

under different scenarios in Figure 5. For comparison, we also include the empirical power and FDR curves of *ℛ*_lasso_ (obtained via the model-X knockoff filter using the lasso coefficient difference statistic).

**Figure 5.**
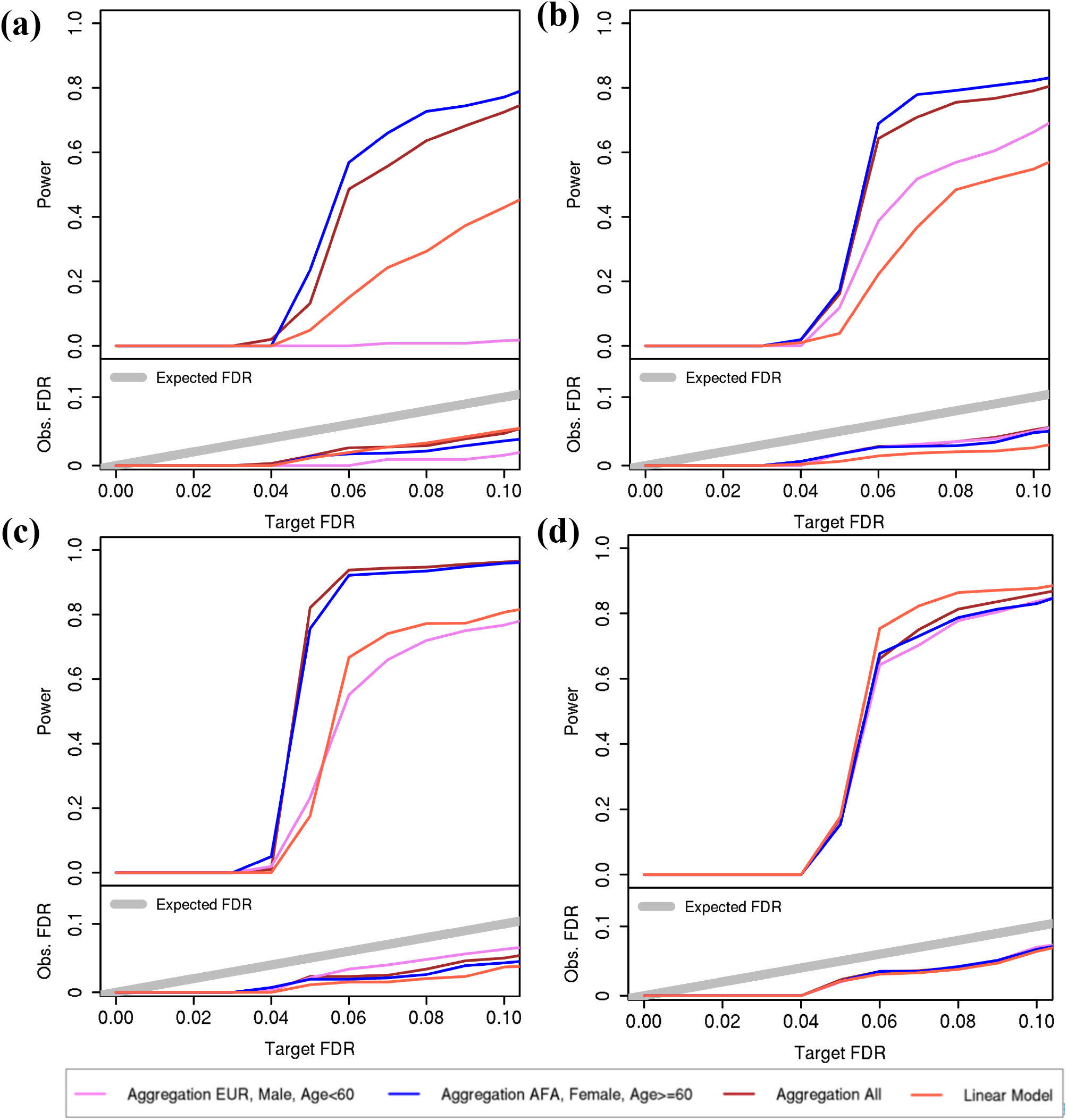
Empirical power and FDR of *ℛ*_population_, obtained by aggregating localized test statistics *W*_***x***,*j*_’s corresponding to various subsets of evaluation points, under different scenarios.

As shown in S1.D, provided that the subgroups are defined prior to observing the data, all these population selection sets are separately equipped with FDR control. In the first three scenarios, *ℛ*_population_ shows superior power to *ℛ*_lasso_, indicating the advantage of aggregating localized evidence against the false *H*_*j*_ *‘*s by accounting for individual heterogeneity. Moreover, *ℛ*_old_afa_women_ demonstrates greater power than *ℛ*_young_eur_men_ in the first three settings, which aligns with Figure 3 where *ℛ*_***x***_’s of old AFA women exhibited the highest power due to strong localized evidence against the false *H*_*j*_ *‘*s, whereas young EUR men had the lowest power. Additionally, the power of *ℛ*_old_afa_women_ is comparable to *ℛ*_population_, suggesting that old AFA women have dominating localized evidence against the false *H*_*j*_’s. In contrast, *ℛ*_young_eur_men_ generally shows lower power than *ℛ*_lasso_ and has zero power in scenario 1, where there is no localized evidence against the false *H*_*j*_ *‘*s for young EUR men. In scenario 4, where no sub-population heterogeneity exists, we observe no significant difference across the power curves.

### F. Simulation results: replicability analysis

In this section, we demonstrate how the strategy proposed in Section 3.E enhances the replicability of our method. Along with assessing *replicability* of our method to return consistent localized selection sets for the same ***X*** = ***x*** across similar training datasets, we also evaluate the *stability* of the method by examining its ability to produce stable selection sets across multiple runs on the same training dataset.

For each training dataset 𝔻 = {𝕏, ***Y***} of size *n* = 1000 generated under Scenario 3 from Section 2.C, we implement our procedure, with and without derandomization, 50 times to compute localized feature selection sets for 1000 evaluation points as in Section 2.D. To measure *replicability*, we compute the average Jaccard similarity of both individual- and population-level selection sets obtained across single runs on the different training datasets (denoted by 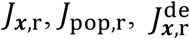, and 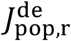 for individual- and population-level, with and without derandomization, respectively). Similarly, we also measure *stability* across multiple runs on the same training datasets (denoted by 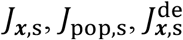, and 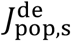). Details on the computation of Jaccard similarities are provided in Section S1.G.

As shown in Figure 6 (a), the proposed *replicability*-enhancing strategy significantly improves the *stability* of selection sets produced by the localized feature selection method. Without derandomization, the *J*_***x***,*s*_ values are below 0.4 for most localized selection sets, indicating considerable variability in inference results due to the randomness of knockoffs generation. In contrast, with *M* = 15 runs and the inclusion-rate-based derandomization strategy introduced in Section 3.E, the 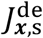 values mostly exceed 0.8, reflecting much higher stability. A similar improvement in stability is observed for the population selection set from *J*_*pop,s*_ = 0.652 to 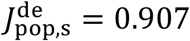, which is higher than most localized selection sets.

**Figure 6.**
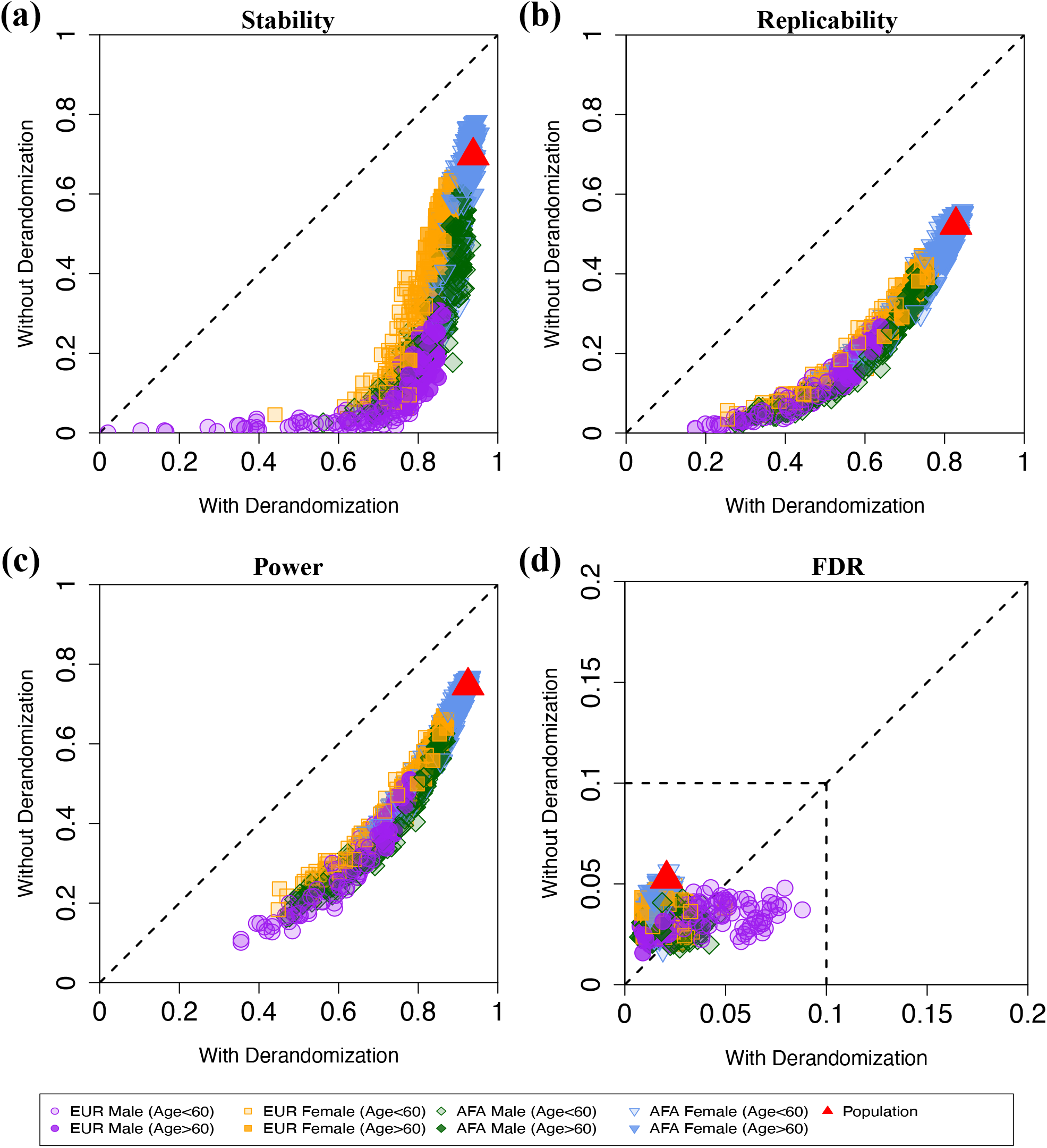
Empirical comparisons between selection sets obtained **with** (horizontal axis) and **without** (vertical axis) derandomization across different evaluation measures under the target FDR level 0.10.

Figure 6 (b) further illustrates the enhanced *replicability* of inference results. Without derandomization, most *J*_***x***,*r*_ values are below 0.2, with none exceeding 0.6. In contrast, nearly all 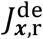 values surpass 0.2, with half exceeding 0.6. The population selection set shows a similar improvement, with increase from *J*_*pop,r*_ = 0.*5*22 to 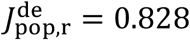, which is higher than most of localized selection sets.

In addition to the improvements in stability and replicability, specific patterns emerge with respect to demographic features. Among the population, localized selection sets for evaluation points with *E*_1_ = *E*_2_ = 1 and a large *E*_3_, which correspond to higher power, are the most stable and replicable. In contrast, localized selection sets for evaluation points with *E*_1_ = *E*_2_ = 0 and a small *E*_3_ exhibit the lowest stability and replicability. This is reasonable as high power of localized selection sets suggests that more important features are consistently selected, leading to both greater stability and replicability.

Finally, Figures 6 (c) and 6 (d) demonstrate that derandomization also improves power at both the individual and population levels while maintaining FDR control at the target level 0.10. This is due to a reduced chance of missing important features across multiple runs, increasing their inclusion in the derandomized selection set.

## 3. Methods

### A. Setting

Consider a dataset 𝔻 = {𝕏 = [***x***_1_, …, ***x***_*n*_]^*T*^, ***Y*** = (*y*_1_, …, *y*_*n*_)^*T*^} that contains *n* individuals, where ***x***_*i*_ = (*x*_*i*1_, …, *x*_*ip*_)^*T*^and *y* represent the *p* features and the response of the *i*-th individual, respectively (*i* = 1, … *n*). The evaluation of the conditional independence hypotheses below is employed for the selection of significant features.

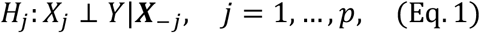

where ***X***_–*j*_ = (*X*_1_, …, *X*_*j*–1_, *X*_*j*+1_, …, *X*_*p*_)^*T*^ records all features except *X*_*j*_. A non-null means that the *j*-th feature has predictive power beyond that afforded by all the other features.

To test these hypotheses, we introduce localized test statistics *W*_***x***,*j*_ in Section 3.C that assess the evidence against each *H*_*j*_, focusing on a neighborhood of the feature space around a given ***x*** =(*x*_1_, …, *x*_*p*_) ^*T*^. We use these to construct a collection of rejection sets *ℛ*_*x*_ such that

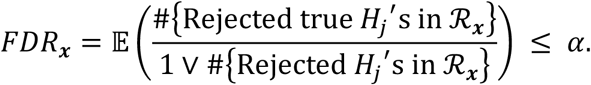

This error rate control ensures that, on average, at least (1 − *α*) × 100*%* of the features discovered in *ℛ*_***x***_ are truly significant.

Aggregating the localized test statistics across all individuals in 𝔻 (see details in Section 3.D), i.e., 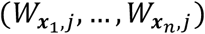 for *j* = 1, …, *p*, yields an overall discovery set *ℛ*_population_ that accounts for heterogeneous signals, such that

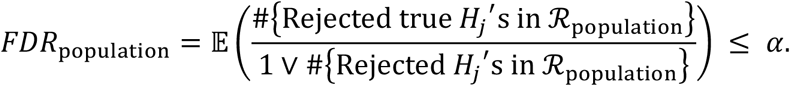

Both the localized and the aggregated test statistics are testing the same conditional independence hypotheses *H*_*j*_. To appreciate their different flavors and provide a prelude to our method, it is useful to think about the many possible departures from the null *H*_*j*_. If *H*_*j*_ holds, then the distribution of *Y*|***X*** remains unchanged regardless of the value of *X*_*j*_, implying that

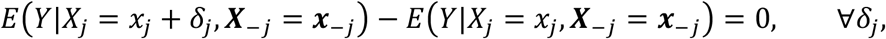

for any given feature values ***x*** = (*x*_*j*_, ***x***_*-j*_)^*T*^. The localized test statistics, which characterize the localized importance of *X*_*j*_ in a neighborhood of ***x***, are tailored to detect localized departures from the null *H*_*j*_ based on an estimate of 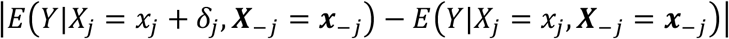.

Since different feature values ***x*** potentially result in different localized selection sets *ℛ*_***x***_, we refer to this as *localized feature selection*, distinguishing it from the more commonly studied *population feature selection* that returns a common subset of features *ℛ*_population_.

### B. Localized feature importance scores

Building on the intuition from the previous section, a natural way of quantifying the localized importance of *X*_*j*_ for the response *Y* in a neighborhood of ***x*** is given by

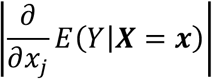

Specifically, this gradient-based measure quantifies how much the expected response would change if the *j*-th feature were perturbed. When (***X***, *Y*) follows a linear model, such a gradient-based measure is a constant throughout the whole population. When (***X***, *Y*) follows a non-linear model, this gradient-based measure is expected to vary across individuals and provide localized information of importance at different feature values. In a genomics example where *Y* denotes an outcome and *X* the expression level of a gene, 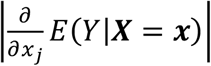 measures the expected rate of change in the outcome when regulating the expression of the gene. This value can be larger for some patients, suggesting that the same increment in the expression level may correspond to different rates of change.

In addition to the gradient, there exist other feature importance scores in the XAI literature that provide localized evidence against the *H*_*j*_’s. Examples include the Shapley value ^40–42^ and the estimated coefficient in localized surrogate models under the LIME framework ^43, 44^. These alternative feature importance scores can also be employed in our method.

### C. Localized feature selection via model-X Knockoffs

#### Construction of knockoffs as synthetic controls

To perform localized feature selection via model-X knockoffs, the first step is to generate knockoffs 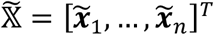 that serve as a form of synthetic controls for the observed features 𝕏 = [***x***_1_, …, ***x***_*n*_]^*T*^. Following the definition in Candès et al. ^11^, 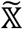 are valid knockoffs of 𝕏 if the following two properties are satisfied: (a) *conditional independence:* 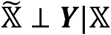, which states that 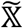 provides no additional information on ***Y***; and (b) *distributional exchangeability:* the joint distribution of 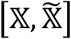 remains unchanged if any pair of columns 𝕏_*j*_ and 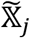 are swapped,

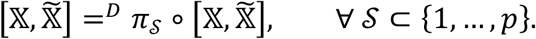

Here, the permutation *π*_*𝒮*_ swaps 𝕏_*j*_ (the *j*-th column of 𝕏) with 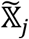 (the *j*-th column of 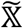) for all *j* ∈ *𝒮*. Specifically, property (a) is satisfied as long as only the covariate matrix 𝕏 is used when generating knockoffs 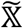. Property (b) implies that, with only the information from 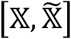, the analyst cannot possibly distinguish 𝕏_*j*_ from 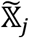. Several approaches have been proposed in the literature for knockoff generation, including second-order knockoffs ^11^, hidden Markov model knockoffs ^45^, deep knockoff machines ^46^, Metropolized knockoff sampler ^47^, and intertwined probabilistic factors decoupling (IPAD) ^25^. In this article, we adopt the IPAD procedure to generate knockoffs with details provided in Section S1.A of the supplementary material.

According to property (b), 𝕏_*j*_ and 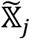 can only be distinguished via their associations with the response ***Y***. Based on this interpretation, we can fit a machine learning model on the extended dataset 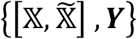 to learn the associations of all 𝕏_*j*_’s and 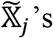 with the response ***Y***. By comparing the importance of 𝕏_*j*_ and 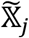, we can select a subset of important features for the response ***Y*** with FDR control. The above reasoning is correct and applies to almost any machine learning model used, as long as the generated knockoffs satisfy both properties (a) and (b) and the fitted machine learning model obeys some mild technical requirements detailed below.

#### Model fitting

Having knockoffs 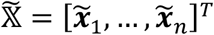, the next step is to fit a machine learning model *h*(⋅) on the extended dataset 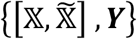 to estimate the conditional expectation 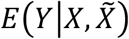.

We then calculate the scores *T*_*j*_’s and *T*_*j*+*p*_’s that measure the importance of *X*_*j*_’s and 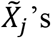 to predict the response *Y*. However, to ensure that *T*_*j*+*p*_ is a negative control for *T*_*j*_ which is indistinguishable from *T*_*j*_ when *H*_*j*_ is true, the algorithm *𝒜* used to obtain the fitted model *ĥ*(⋅) needs to be equivariant to any swapping operation *π*_*𝒮*_ applied on 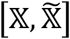. That is to say, 𝕏_*j*_ and 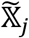 have the same role in the algorithm *𝒜*, no matter whether the algorithm *𝒜* is deterministic or stochastic. The formal definition of this algorithmic symmetry is provided in Section S1.C of the supplementary material. Examples of model fitting methods *𝒜* satisfying this algorithmic symmetry property include fitting generalized linear models, random forests, neural networks in which 𝕏_*j*_ and 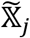 are treated symmetrically in the input layer. In Sections 2.B and 2.C, we employ a quadratic regression model detailed in Section S1.B of the supplementary material. We note that our procedure provably controls the FDR at the desired level regardless of the correctness of the fitted model, although a good model generally leads to higher power.

#### Localized feature selection

We follow the line of thinking in Section 3.B and use the gradient of 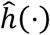 at ***x*** to assess the localized evidence against *H*_*j*_ for an individual with features ***X*** = ***x***. For any differentiable fitted model 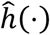, we use the absolute values of derivatives 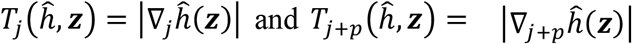 to calculate the localized importance of *X*_*j*_ and 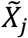 for the response *Y*, evaluated at ***z*** = (***x, x***). A detailed discussion about the choice of evaluation data value ***z*** can be found in Supplementary Materials S1.C and S1.D. We then combine 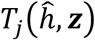 and 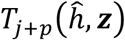 into a localized test statistic

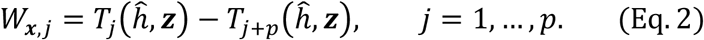

Intuitively, as knockoffs 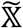 are generated without using the information from 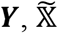 provides no additional information about *Y* beyond 𝕏. Therefore, the gradient 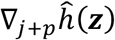 is expected to be small, while 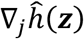 is expected to be around 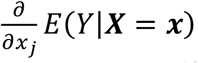. As a result, if there is strong evidence against *H*_*j*_ at ***x***, we expect 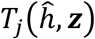 to dominate 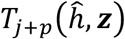 and, therefore, *W*_***x***,*j*_ to be large and positive; if *H*_*j*_ is null, then we expect 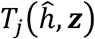 and 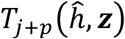 to be similar. Formally, we show in Section S1.D of the supplementary material that the *W*_***x***,*j*_’s satisfy the coin-flip property (i.e., signs of *W*_***x***,*j*_’s for those true *H*_*j*_ are i.i.d. coin flips conditional on all 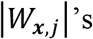 and signs of *W*_***x***,*j*_’s for those false *H*_*j*_’s ^11^). We apply the knockoff filter ^11^ on *W*_***x***,*j*_’s to obtain a subset *ℛ*_***x***_ of features that have strong evidence against *H*_*j*_’s at ***x***. In short, the procedure to obtain *ℛ*_***x***_ is summarized as follows.

Using the model 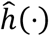 fitted on the entire data and knockoffs,

1. Compute the localized importance score and its knockoff 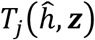, and 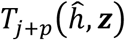 at ***z*** = (***x, x***)(*j* = 1, …, *p*);
2. Combine 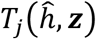 and 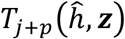 into a localized test statistic 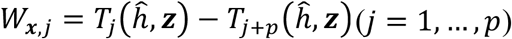;
3. Under the target FDR level *α* ∈ (0,1), apply the knockoff filter ^11^ on *W*_***x***,*j*_’s to obtain a subset of features *ℛ*_***x***_ that have strong evidence against *H*_*j*_’s at ***x***,

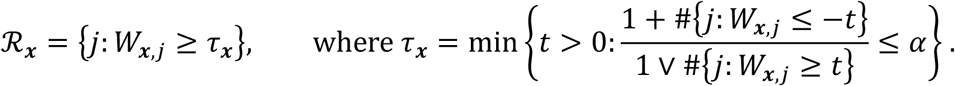

As the gradient 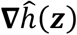) generally varies across different ***z*** = (***x, x***)*’*s, implementing the above procedure for different individuals selects different subsets of features, all obeying FDR control at level *α*, see Section S1.D of the supplementary material.

*Remark: Here, the FDR control of ℛ*_***x***_ *applies to the hypotheses* 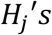, *which indicate whether each feature X*_*j*_ *has any (conditional) influence on the response Y in the population. The localized test statistics* 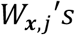 *provide a localized ranking of features based on their impact on Y in the neighborhood of* ***X*** = ***x*** *within the sample space. Since ℛ*_***x***_ *generally varies with* ***x***, *it offers insights how the influence of important features on the response differs across regions of the input space*.

In addition to the selection set *ℛ*_***x***_, assigning priority scores to the features within *ℛ*_***x***_ is also crucial for real-world applications. For example, in designing personalized therapies, it is impractical to simultaneously target all important biomarkers for an individual due to safety or cost constraints. Instead, practitioners typically prioritize biomarkers in a decreasing order of importance. Motivated by this, we propose the augmented selection vector for a generic individual with feature values ***X*** = ***x***:

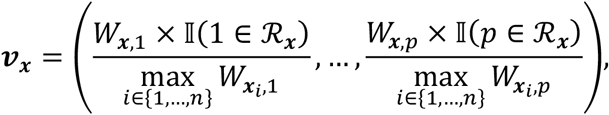

where each ***x***_*i*_ is a vector of covariates from the sample. Here, the *j*-th component of ***v***_***x***_ is nonzero if and only if *j* ∈ *ℛ*_***x***_, and its value corresponds to the localized test statistic *W*_***x***,*j*_ normalized by the maximum across the entire sample. The normalization ensures that no single feature dominates others in subsequent unsupervised analysis tasks. For a set of candidate feature values of interest 𝔻_*eval*_, we compute the augmented selection matrix 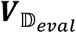, formed by concatenating the augmented selection vectors ***v***_***x***_’s for all ***x*** ∈ 𝔻_*eval*_. This matrix can then be used in downstream exploratory analyses to uncover hidden structure in the data.

#### D. Aggregating localized feature selection for population-level inference

Our framework can also be used in performing powerful *population feature selection* by aggregating localized inference results. For example, aggregating 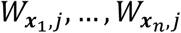 as

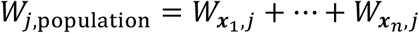

can provide an overall measure of evidence against *H*_*j*_. The selection set *ℛ*_population_ at the population level under the target FDR level *α*_agg_ ∈ (0,1) can be obtained by applying the knockoff filter ^11^ to *W*_1,population_, …, *W*_*p*,population_:

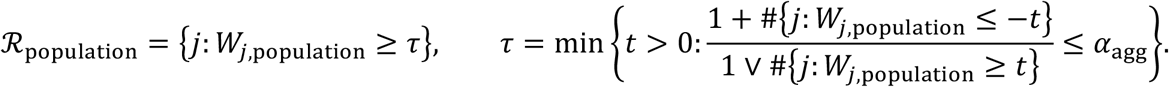

We show in Section S1.E of the supplementary material that the *W*_*j*,population_*’*s obey the coin-flip property and therefore *ℛ*_population_ controls the FDR at level *α*_agg_. As the localized test statistic *W*_***x***,*j*_ measures the strength of evidence against *H*_*j*_ at ***x***, the localized feature importance aggregation can capture potential heterogeneous effects, thereby improving statistical power for selecting important features at the population level. In Section S1.E of the supplementary material, several other aggregation methods, including e-value-based aggregation ^48^ and inclusion-rate-based aggregation ^49^, are introduced. We recommend using the simple sum as the aggregation function, as it is the most straightforward and empirically powerful option, as demonstrated in Section S2.A.

Finally, we remark that localized feature importance aggregation can also be done at the level of subpopulations that are predefined prior to looking at the data. For example, if a study is interested in health disparity and would like to evaluate heterogeneous effects based on sex, one could aggregate the localized test statistics according to cohorts defined by sex and then apply the knockoff filter respectively.

#### E. Strategy to improve replicability

Our proposed method enables both localized and population-level feature selection with provable FDR control, ensuring enhanced replicability over existing methods, which generally lack control over the number or proportion of false positives. In this section, we provide an additional strategy to further improve the replicability of our method.

According to National Academies of Sciences, Engineering, and Medicine et al. ^50^ and Nichols el al. ^51^, replicability refers to the ability to draw consistent conclusions from replicate studies that use the same inference procedure and address the same scientific question on similar datasets. In the context of localized feature selection, we define replicability as

- For any target individual with features ***X*** = ***x***, implementing our *localized feature selection* on similar training datasets generated from the same population would return similar selection sets *ℛ*_***x***_’s.

The *replicability* of the proposed method in returning selection sets *ℛ*_***x***_’s is influenced by two factors: (a) the sampling variation of the training dataset 𝔻 = {𝕏, ***Y***}, and (b) the *stability* of our approach, characterized by the randomness in knockoff generation and model fitting (if the fitting algorithm *𝒜* employed is stochastic). While the sampling variation is unavoidable, one can enhance the stability so that repeated runs of our method on the same training dataset yield stable inference results. To achieve this, we propose a derandomization strategy that performs multiple runs of our localized feature selection procedure and aggregates the results from these runs. Specifically, for a target candidate feature value ***X*** = ***x***, we obtain *M* selection sets 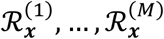 under the FDR level *α* by performing *M* independent runs of our procedure. We then compute the derandomized selection set 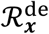 by applying the inclusion-rate-based procedure ^49^ to 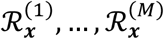 with details in Section S1.F of the supplementary material. The derandomized selection set 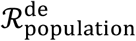 at the population level can be also obtained in an analogous manner. As shown by the experimental results in Section 2.F, adopting the derandomization strategy improves the *stability* of our method with decreased variation in the selection sets. It also improves the *replicability* at both individual and population levels, while *empirically* controlling the FDR at the desired level.

Analogously to Section 3.C, we define the augmented selection vector 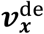 corresponding to the derandomized selection set 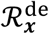 as

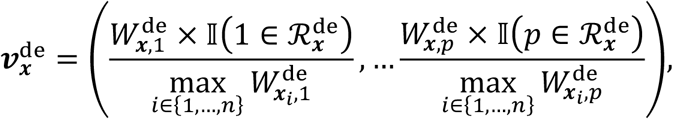

where 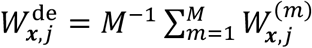 is the average of the localized importance scores 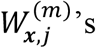 among *M* runs of our procedure.

## 4. Discussion

In this paper, we introduced a novel framework for localized feature selection with false discovery rate control by leveraging model-X knockoffs. Our framework uses localized feature statistics *W*_***x***,*j*_ to assess the strength of localized evidence against the conditional independence hypotheses *H*_*j*_ across diverse individuals, yielding heterogeneous localized selection sets *ℛ*_***x***_’s. By aggregating the *W*_***x***,*j*_’s across individuals, the framework combines heterogeneous localized evidence against false *H*_*j*_’s to produce a powerful population-level feature selection set with FDR control. Below, we outline several potential directions for future research.

First, it is important to note that while our localized feature selection procedure effectively identifies localized rejection sets for different values of the covariates and captures effect heterogeneity among important features, we are still testing the conditional independence hypotheses *H*_*j*_ at the population level. Specifically, the FDR control for *ℛ*_***x***_ established in this paper is with respect to the population-level hypotheses. This guarantees that most discoveries correspond to truly important variables for the population, which is desirable since the set of important variables at the population level typically encompasses those relevant to specific subgroups. However, our approach does not necessarily ensure error-rate control for localized conditional independence hypotheses, as defined in works such as Sesia and Sun ^52^ and Gablenz et al. ^53^. While in applications where localized subgroups and the population share the same important features but with potentially different effect sizes, controlling FDR for population-level conditional independence hypotheses also implies FDR control for localized conditional independence hypotheses, it would be an interesting future work to extend our method within the theoretical framework of Sesia and Sun ^52^ and Gablenz et al. ^53^.

Second, the past few decades have witnessed a dramatic surge in the volume and variety of health-related digital information. Beyond electronic clinical data, which capture demographic characteristics (e.g., gender, age) and clinical records (e.g., pathological and physiological history), next-generation sequencing technologies, along with advances in imaging- and fluid-based biomarker approaches {Thal, 2025 #1450}, have generated a wealth of omics data, enabling highly detailed descriptions of individuals ^54–56^. Future work could apply our framework to infer insights from integrated health information spanning multiple data types. Jointly analyzing the diverse omics data of individuals, potentially from hidden subpopulations, may offer new insights into the heterogeneous biological pathways associated with traits of interest.

Third, while our framework employed a lasso-penalized quadratic regression model, other nonlinear modeling approaches could be explored. For example, developing appropriate deep learning models to capture complex relationships between features and the response might enhance power, particularly for large datasets. Note that the critical requirement of our framework is the symmetric treatment of original variables and their knockoffs. Therefore, in practice, cross-validation can be used to select among multiple models with algorithmic symmetry for estimating *h* without compromising FDR control.

Finally, our framework also raises new research questions beyond localized feature selection. For instance, investigating discoveries with heterogeneous effects across individuals, as revealed by the augmented selection matrix, and probing their underlying biological mechanisms through downstream analyses could provide valuable insights. Such inquiries could advance personalized medicine by informing the development of tailored drugs and therapies. Along these lines, the development of interventions that differentially affect newly identified features would make possible a modeling to experimental testing feedback loop. The detection of previously unrecognized, yet robust, distinct features across a given cell population or subcellular entity, will further advance precision approaches, contribute to cellular context molecular mechanism elucidation and will help to make possible the detection of therapeutic effects that might otherwise be overlooked.

## Data Availability

The single-cell RNA sequencing (scRNA-seq) dataset analyzed in this study is publicly available from the NCBI GEO database under accession number GSE163577.

## Code Availability

The proposed method has been implemented as an efficient R package, available at https://github.com/circustata/LocalizedFeatureSelection.

## Acknowledgments

Z.C. was supported by the Simons Foundation under award 814641. C.S. was supported by the grants NIH R56HG010812 and NSF DMS2210392. E.J.C. was supported by the Office of Naval Research grant N00014-20-1-2157. Z.H. was supported by NIH/NIA award AG089509, AG066206 and AG066515.

## Supplementary Materials

### S1. Supplementary details of the inference procedure

#### A. The IPAD procedure for knockoffs generation

In this article, we utilize the IPAD procedure from Fan et al. ^25^ to generate knockoffs. According to Candès et al. ^11^, knockoffs 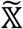 satisfies both the conditional independence 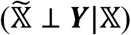 and the distributional exchangeability 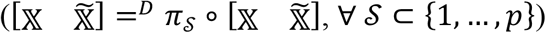. If observed features 𝕏 follow the factor model

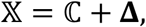

where ℂ = 𝔽𝕃^*T*^, 𝔽 = [***f***_1_, …, ***f***_*n*_]^*T*^ ∈ *ℛ*^*n*×*r*^ is the random matrix of unobserved factors, 𝕃 = (***L***_1_, …, ***L***_*p*_)^*T*^ ∈ *ℛ*^*p*×*r*^ is the deterministic loading matrix, and **Δ** = [***δ***, …, ***δ***_*n*_]^*T*^ ∈ *ℛ*^*n*×*p*^ is the matrix of random errors drawn i.i.d. from some parametric models *g*_*η*_(⋅) independent to 𝔽, then knockoffs 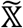 can be generated as

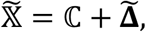

where 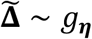 is a random error matrix independent of both 𝔽 and **Δ**.

In practice, however, the matrix ℂ and parameters ***η*** are unobserved. Therefore, we generate knockoffs 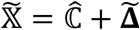 with the estimated 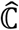 and 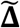 generated from 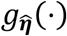 using the estimated 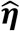. In this article, by assuming that ***δ***_1_, …, ***δ***_*n*_ are i.i.d. multivariate Gaussian errors with zero mean and covariance matrix *diag*(*η*_1_, …, *η*_*p*_), we use the principal component (PC) estimators 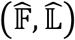to obtain the estimated 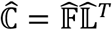 and the estimated 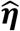 by

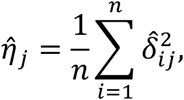

where 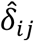 is the (*i, j*)-entry of 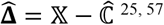.

In our real data application, we use hyperparameter *r* = 271, where *r* is the number of PCs used for knockoffs construction. The selection of *r* is based on examining the off-diagonal elements of the covariance matrix of the residuals to ensure valid knockoffs construction. Specifically, we iteratively increase *r* and calculate the off-diagonal elements of the residual covariance matrix until their average value falls below the predefined threshold 0.01. To check whether the IPAD procedure generates approximately valid knockoffs, we compare the empirical correlations of *X*_*j*_ and *X*_*k*_ (*j ≠ k*) with the empirical correlations of 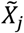and 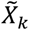 (left panel of Supplementary Figure 5) and compare the empirical correlations of *X*_*j*_ and *X*_*k*_ with the empirical correlations of *X*_*j*_ and 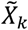 (right panel of Supplementary Figure 5) over all pairs of 1000 randomly sampled features (or 1000 randomly sampled feature pairs). We observe that for most of the feature pairs, the empirical correlations of 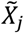 and 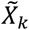(and of *X*_*j*_ and 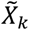) match those of *X*_*j*_ and *X*_*k*_, indicating that the generated knockoffs approximately satisfy the *distributional exchangeability* property required for knockoff validity (11).

#### B. Details of fitting the quadratic regression model that retains knockoffs exchangeability

The model-fitting method *𝒜* used in Section 2 is lasso-penalized regression on the quadratic model,

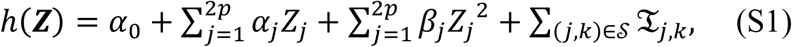

where

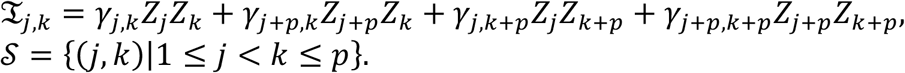

Here, *Z*_*j*_ corresponds to *X*_*j*_, and *Z*_*j*+*p*_ corresponds to 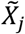. The localized test statistic *W*_***x***,*j*_ is computed as

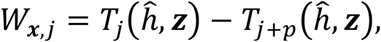

where

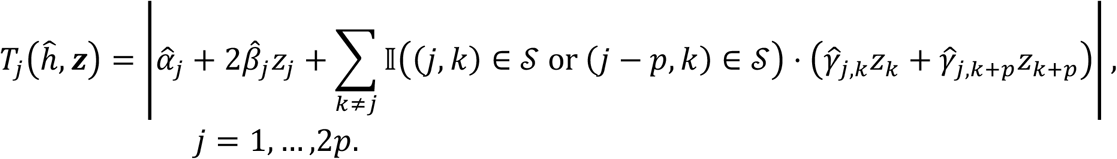

When *h*(⋅) is a linear model, *W*_***x***,*j*_ in (*Eq*. 2) reduces to the lasso coefficient-difference (LCD) statistic used in Candès et al. ^11^, which is constant over different ***x***’s.

Since directly fitting the full model *h*(⋅), which includes 2*p*^2^ + 2*p* + 1 parameters, is computationally expensive and risks overparameterization, we adopt the strategy of Sesia and Sun (2022). Specifically, we propose a knockoff-invariant screening method (Algorithm S1) to enhance the model fitting performance and the power of localized feature selection.

The knockoffs invariant screening method performs screening in three steps. First, by fitting the initial additive model with only linear and quadratic terms without interactions, we select *N*_1_ features that are most likely to have effects on the response (steps 1-4). We then fit the extended model by adding all interaction terms among the *N*_1_ most promising features (step 5). Finally, we refine the model fitting by pruning interactions with neglectable effects to achieve the lowest BIC (steps 6). Given the refined model with linear and quadratic terms and selected interactions, we obtain the final fitted model 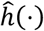 via lasso regression with cross-validation.

##### Algorithm S1 Knockoffs invariant screening.

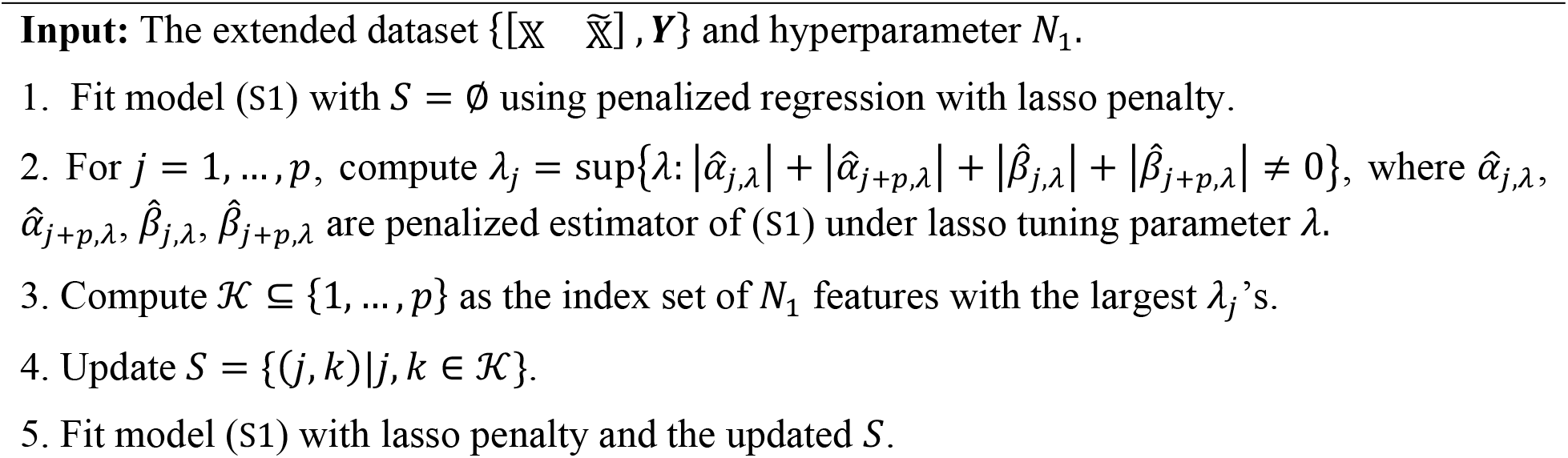
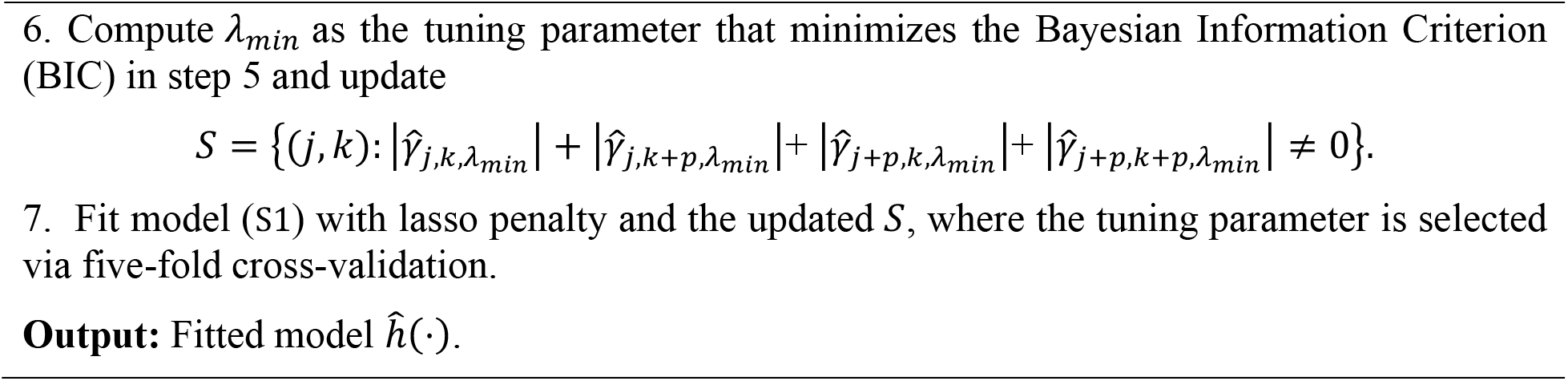

It is straightforward to show that Algorithm S1 is equivariant to the swapping operation *π*_*𝒮*_ applied to 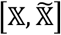. Thus, Algorithm S1 maintains the algorithmic symmetry property required for valid FDR control while reducing model complexity via multi-stage screening. Although Algorithm S1 is proposed for the fitting model (S1), the screening strategy can be adapted to regularize other machine learning models with the algorithmic symmetry property.

In the following, we use Scenario 3 from Section 2.C as an example to illustrate how the knockoffs invariant screening strategy enhances our method’s performance in terms of improved statistical power and computational efficiency. For comparison, in addition to implementing the knockoff invariant screening procedural defined in Algorithm S1, we also implement the following variations: (a) the directly fitted quadratic model (S1) without knockoffs screening, and (b) the fitted quadratic model (S1) with *S* computed in step 4 of Algorithm S1.

As shown in Supplementary Figure 1(a), the power associated with the localized selection sets is the highest when using the fitted quadratic model (S1) with *S* computed by Algorithm S1, whereas the power is lowest when no screening is performed. Furthermore, Supplementary Figure 1(b) demonstrates that computation time decreases as more stages of screening are incorporated. Thus, both stages of knockoffs invariant screening in Algorithm S1 (steps 1-4 and steps 5-6) improve the power and computational efficiency of localized feature selection, highlighting the effectiveness of incorporating knockoffs invariant screening.

**Supplementary Figure 1.**
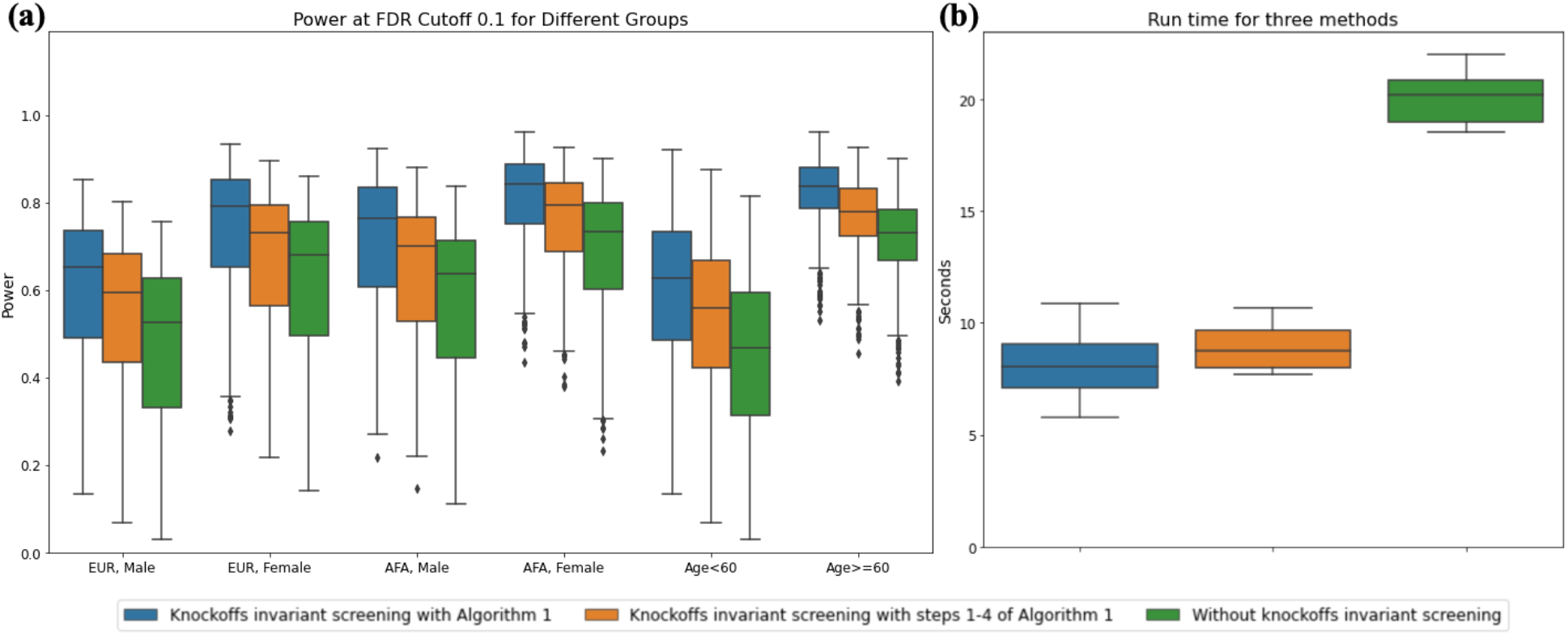
Comparison of different methods using fitted quadratic models under Scenario 3 from Section 2.C. (a) Comparison of the statistical power of *ℛ*_***x***_ *‘*s across diverse profiles of evaluation points, using the knockoff filter with a target FDR level of *α* = 0.10 and different fitted quadratic models under Scenario 3. (b) Comparison of run times for the different fitted quadratic models (CPU: Apple M1 Pro, 3.22 GHz; RAM: 16GB).

#### C. Technical conditions required for the inference procedure

As described in Section 3, there are five main steps to obtain a subset of features *ℛ*_***x***_ with strong evidence against *H*_*j*_’s around ***X*** = ***x*** from the dataset 𝔻 = {𝕏, ***Y***}. These steps include: generating knockoffs 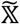, using an algorithm *𝒜* to obtain a fitted model 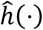, extracting localized importance scores 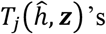 and 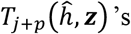 at ***z*** = (***x, x***), computing localized test statistics *W*_***x***,*j*_ *‘*s by combining 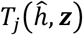 and 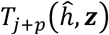, and applying the knockoff filter ^11^ to *W*_***x***,*j*_’s. In addition to the validity of the knockoffs 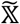, some properties are required for the algorithm, localized importance scores, and localized test statistics as detailed below.

1. **Algorithmic Symmetry**: As stated in Section 3.C, the algorithm *𝒜* used to obtain a fitted model 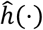 must be equivariant to any swapping operation *π*_*𝒮*_ applied to 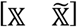. Specifically,
  a. if *𝒜* is a deterministic algorithm, applying *𝒜* to the swapped dataset 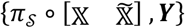 should yield the model 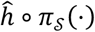 such that

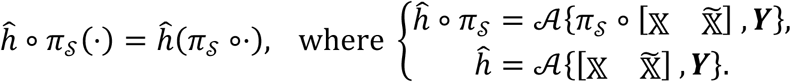
  b. if *𝒜* is a stochastic algorithm, applying *𝒜* to the swapped dataset 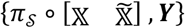 should yield the model 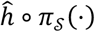 such that

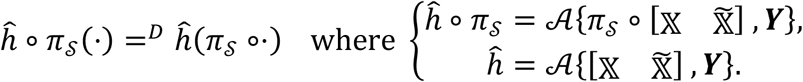 When *𝒜* is a deterministic algorithm, algorithmic symmetry implies that applying algorithm *𝒜* to the swapped dataset 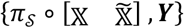 is equivalent to swapping the corresponding arguments of the fitted model 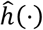, leading to the model 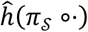.
2. **Feature Score Symmetry**: Feature importance scores extracted from the fitted model 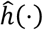 need to be symmetric with respect to the swapping operation *π*_*𝒮*_. Specifically, this is expressed as

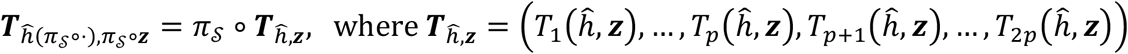

for any ***z*** ∈ *ℝ*^2*p*^ and any *S* ⊂ {1, …, *p*}. When the fitted model 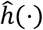 is a differentiable function (e.g., a linear model, quadratic model or neural network with smooth activation functions), it is straightforward that the absolute values of the gradients 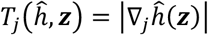 and 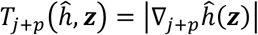 possess the feature score symmetry property because

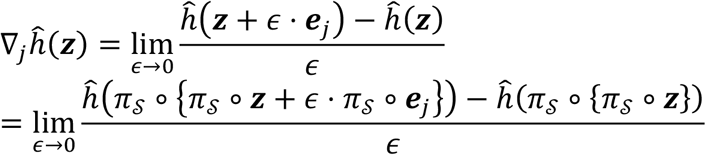

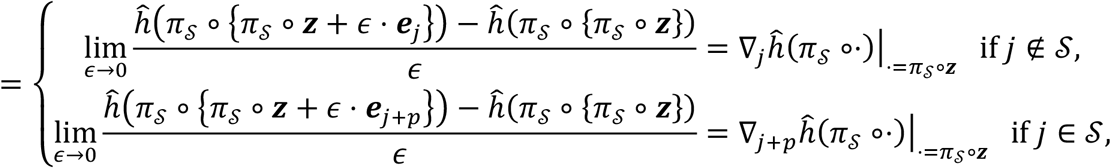

where ***e***_*j*_ and ***e***_*j*+*p*_ are the *j*-th and (*j* + *p*)-th vector of the canonical basis of *ℛ*^2*p*^. However, when 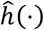 is not a differentiable function (e.g., 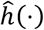 obtained from a random forest), we can no longer construct importance scores and localized test statistics based on gradients. In such cases, feature importance scores can be constructed to measure the localized variation of 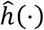 in a manner analogous to gradients. One example is

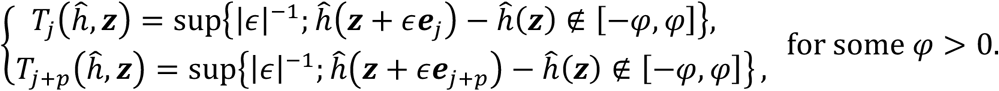 Analogously, it can show that the feature importance score defined above satisfies the required symmetry.
3. **Anti-symmetry of Test Statistics**: To produce the final localized test statistics ***W***_***x***_ = (*W*_***x***,1_, …, *W*_***x***,*p*_), we aggregate the feature importance scores 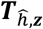 such that

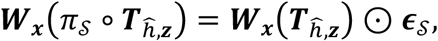

where ***ϵ***_*S*_ is a vector with all entries equal to 1, except for the entries in *S* which are equal to −1, and ⊙ represents the element-wise product. As stated in Candès et al. ^11^, such ***W***_***x***_ can be constructed using

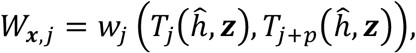

where *w*_*j*_ is an anti-symmetric function such that *w*_*j*_(*a, b*) = *−w*_*j*_(*b, a*) for any *a* and *b*. In addition to 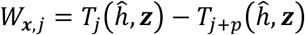, other examples of localized test statistics that are anti-symmetric with respect to 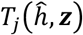 and 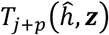 include:

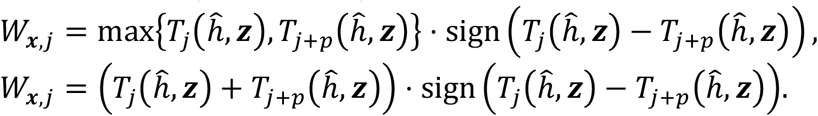

#### D. Proof of FDR control

We show that the *ℛ*_***x***_ obtained in the proposed localized feature selection framework controls the FDR with respect to *H*_*j*_’s under the following four conditions:

1. Validity of knockoffs 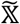:

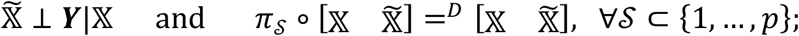
2. Algorithmic Symmetry of the algorithm *𝒜* that produces the fitted model 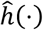;
3. Feature Score Symmetry of 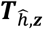;
4. Anti-symmetry of Test Statistics ***W***_***x***_ = (*W*_***x***,1_, …, *W*_***x***,*p*_):

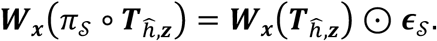

According to the proof of Theorem 3.4 in Candès et al. (11) and Theorems 1 and 2 in Barber and Candès (2015), a sufficient condition for controlling the FDR associated with *ℛ*_***x***_ is that the test statistics *W*_***x***,*j*_’s satisfy the coin-flip property ^11^,

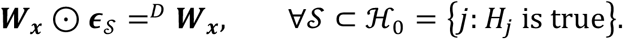

Therefore, to show that *ℛ*_***x***_ controls the FDR, it suffices to prove that the associated ***W***_***x***_ satisfies the coin-flip property, which we establish in Theorem S1 below.

##### Theorem S1.

*Under the four aforementioned conditions (knockoffs validity, algorithmic symmetry, feature score symmetry, and test statistic anti-symmetry), the feature statistics W*_***x***,*j*_’*s obtained from* 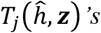 *and* 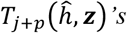 *with* ***z*** = (***x, x***) *possess the coin-flip property if* ***x*** *is either a fixed point or a random point independent of the extended dataset* 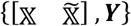.

**Proof of Theorem S1**. By Lemma 2 of Candès et al. ^11^, for any 𝒮 ⊂ *ℋ*_0_,

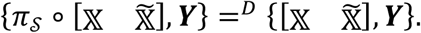

Therefore,

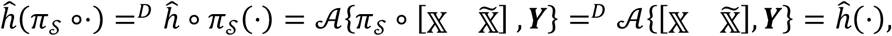

where we used the algorithmic symmetry in the first equality.

Since ***z*** = (***x, x***), we have *π*_*𝒮*_ *∘* ***z*** = ***z*** for any 𝒮 ⊂ *ℋ*_0_. By the independence between ***x*** and 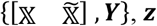 is independent of 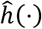 and 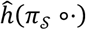. Thus,

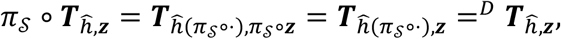

where the first equality follows from the feature score symmetry of 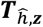.

As a result, the coin-flip property of *W*_***x***,*j*_’s follows by

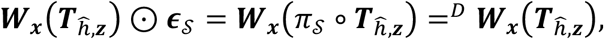

where the first equality is due to the anti-symmetry of the test statistics.

This concludes the proof. However, for some technical reasons, Theorem S1 does not hold when ***x*** is a point in the training dataset A. In the following Theorem S2, we present a modification of Theorem S1 such that the resulting rejection set controls the FDR when ***x*** is a point in A (without loss of generality, we only consider the case where ***x*** = ***x***_1_).

##### Theorem S2.

*With the four aforementioned conditions (knockoffs validity, algorithmic symmetry, feature score symmetry, and test statistics anti-symmetry), the feature statistics* 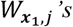 *obtained from* 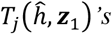 *and* 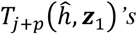 *with* 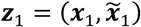 *possess the coin-flip property*.

**Proof of Theorem S2**. By Lemma 2 of Candès et al. ^11^, for any 𝒮 ⊂ *ℋ*_0_, we have

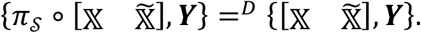

With 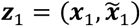, it follows that

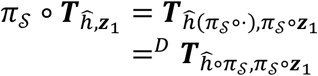

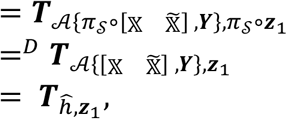

where the first equality follows from the feature score symmetry of 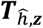 and the second equality follows from the algorithmic symmetry.

Therefore, the coin-flip property of 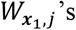 can be proved by

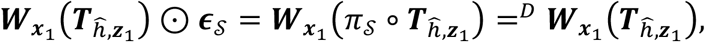

where the first equality is due to the anti-symmetry of the test statistics. This concludes the proof.

Although the FDR control of *ℛ*_***x***_ with ***z*** = (***x, x***) is not provable when ***x*** is a point in A, simulation results suggest that doing so does not cause FDR inflation empirically. Thus, in practice, we suggest using our unified steps in Section 3.C with ***z*** = (***x, x***) for any ***x***, whether an independent data point or one in the training data.

#### E. More details of aggregation approaches for population-level inference

From the inference procedure described in Sections 3.C through 3.D, two types of localized information about the null hypotheses *H*_*j*_’s are generated:

1. **localized test statistics** 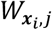 for *j* = 1, …, *p* and *i* = 1, …, *n*.
2. **localized selection sets** 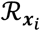 for *i* = 1, …, *n*, each of which has controlled FDR with respect to the null hypotheses 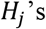.

In this section, we detail how to aggregate either 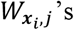 or 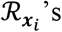 to obtain a population-level selection set *ℛ*_population_ with FDR control with respect to *H*_*j*_’s.

##### Path 1: Aggregating localized test statistics

In this paper, following the procedure outlined in Section 3.D, we aggregate the localized test statistics by summing them across all individuals

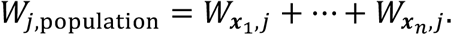

This approach is a specific instance of a broader class of aggregation methods that can be used to obtain population-level feature statistics. In the following Theorem S3, we present a general condition under which the aggregated feature statistics *W*_*j*,population_*’*s satisfy the coin-flip property, which is sufficient for maintaining FDR control.

###### Theorem S3.

*With the four aforementioned conditions (knockoffs validity, algorithmic symmetry, feature score symmetry, and test statistics antisymmetry), if the aggregated feature statistic*

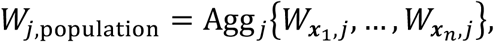

*is obtained via an odd symmetric aggregation rule* Agg_*j*_ *where*

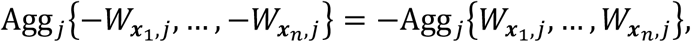

*for j* = 1, …, *p, then the aggregated feature statistics W*_*j*,population_*’s possess the coin-flip property*.

**Proof of Theorem S3**. Since 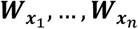 are all obtained from the same extended dataset 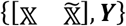, analogous to the proof of Theorem S2, we have

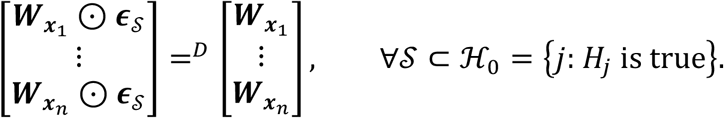

Given the odd symmetry property of the aggregation rules Agg_*j*_’s, we have

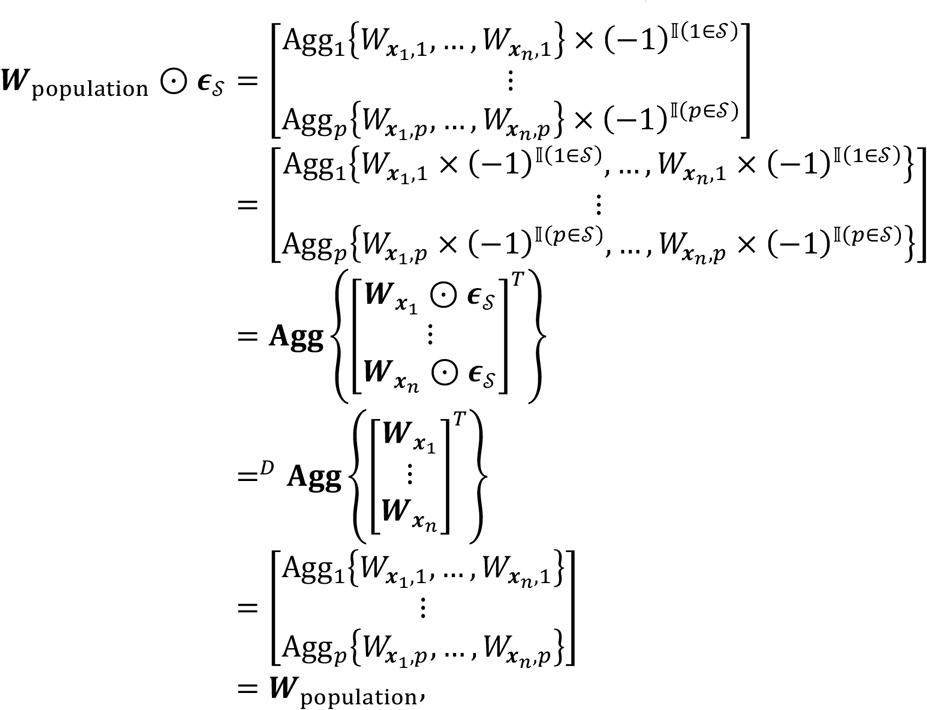

where ***W***_population_ = (*W*_1,population_, …, *W*_*p*,population_)^*T*^. Thus, the coin-flip property of *W*^*j,population*^*’*s holds.

It is evident that summation is an odd symmetric aggregation rule and thus the resulting selection set *ℛ*_population_ has controlled FDR with respect to *H*_*j*_’s. In addition to the summation, examples of other odd symmetric aggregation rules include:

- Weighted sum

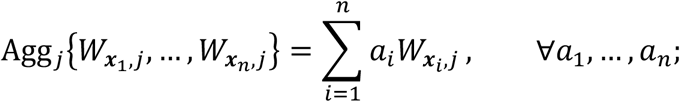
- Sum (or integral) of symmetric quantiles

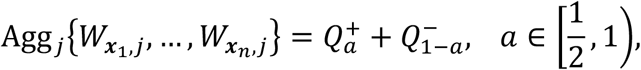

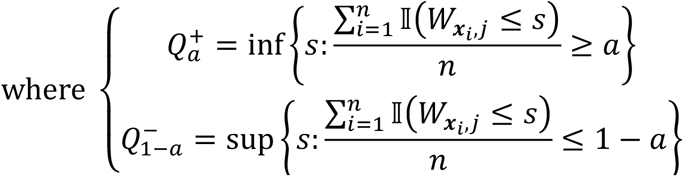
- Sample skewness.

##### Path 2: Aggregating localized selection sets

As the localized selection set 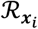 provides a set of *H*_*j*_’s which are believed to be false with strong localized evidence at ***x***_*i*_, aggregating 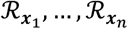 provides a selection set *ℛ*_population_ of *H*_*j*_’s with strong global evidence against the nulls. The e-value aggregation technique ^48^ and the inclusion-rate-based procedure ^49^ are two possible aggregation methods in the literature.

#### E-value aggregation technique

E-values are a recent development in the field of multiple testing, introduced by Vovk and Wang ^58^, and have found extensive use due to their unique properties. Specifically, an e-value with respect to a hypothesis is a random variable with an expectation not larger than 1 when the hypothesis is true. A large e-value implies strong evidence against the corresponding hypothesis. Based on such an interpretation, we employ the e-value aggregation technique ^48^ to obtain the selection set *ℛ*_population_ at the population level using the following steps:

1. **Compute localized e-values**: For *i* = 1, …, *n*, compute localized e-values corresponding to the hypotheses *H*_*j*_ *‘*s from the localized selection set 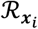 with a target FDR level *α*:

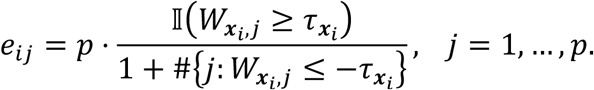
2. **Compute population e-values:** Take the average of the localized e-values to obtain population-level e-values:

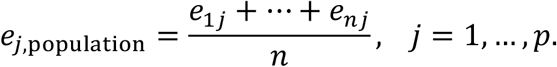
3. **Apply the e-BH procedure:** Implement the e-BH procedure (Wang and Ramdas, 2022; Ren & Barber, 2023) to obtain the selection set *ℛ*_population_ at the population level with respect to the FDR level *α*_agg_ ∈ (0,1)

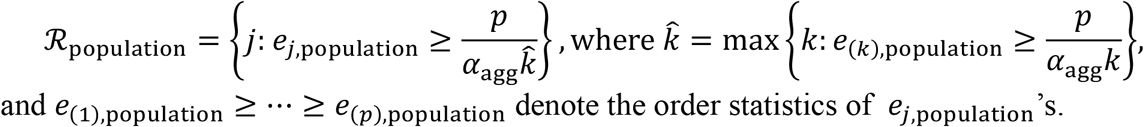

Since for any fixed *i*, Ren and Barber ^48^ showed that *e*_*ij*_’s are valid (generalized) e-values, it is easy to see that *e*_*j*,population_*’*s are also valid (generalized) e-values. Thus, the FDR control (at level *α*_agg_) of the selection set *ℛ*_population_ obtained via the e-BH procedure is guaranteed by Theorem 3 of Wang and Ramdas ^59^. Further, following the recommendation of Ren and Barber ^48^, we use 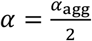 in our experiments.

#### Inclusion-rate-based procedure

The inclusion-rate-based procedure, as introduced by Dai et al. ^49^, provides a method for aggregating localized selection sets *ℛ*_***x***_*i ‘*s in a manner similar to the e-value aggregation technique proposed by Ren and Barber ^48^. The procedure operates as follows:

1. **Compute localized inclusion rates**: For *i* = 1, …, *n*, compute the localized inclusion rates corresponding to the hypotheses *H*_*j*_ *‘*s from the localized selection set 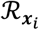:

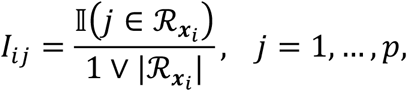

where 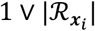 denotes the maximum of 1 and the cardinality of 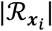.
2. **Compute population inclusion rates:** Aggregate the localized inclusion rates to obtain population-level inclusion rates:

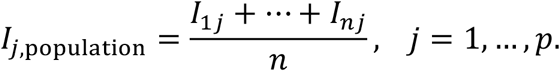
3. **Obtain the population selection set**: Construct the selection set *ℛ*_population_ at the population level with respect to the FDR level *α*_agg_ ∈ (0,1):

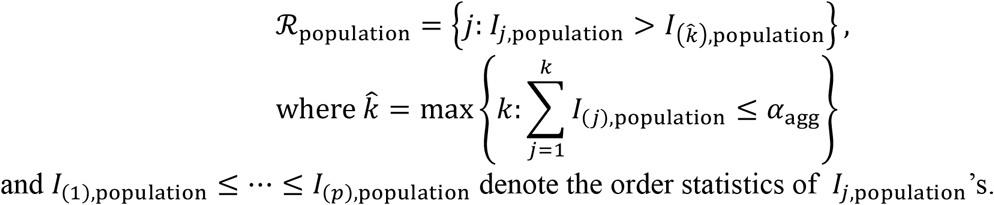

Unlike the e-value aggregation technique, the inclusion-rate-based procedure requires *α* = *α*_agg_. The inclusion rates *I*_*j*,population_*’*s serve as a measure of the relative importance of features *X*_*j*_’s to the response *Y*, based on the following two informal statements:

- The expected contribution of irrelevant features *X*_*j*_ *‘*s (for which *H*_*j*_’s are true) to the sum of localized inclusion rates is bounded by the target FDR level *α* = *α*_agg_,

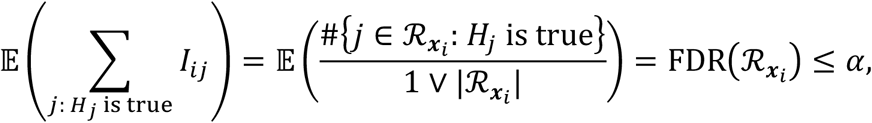

by Theorem S1. Consequently, the average contribution of irrelevant features to the sum of population inclusion rates is also bounded by the target FDR level *α* = *α*_agg_.
- If the feature *X*_*j*_ is selected less (or more) frequently in localized selection sets, it is less (or more) likely to be an important feature.

Intuitively, important features *X*_*j*_ *‘*s with substantial evidence against *H*_*j*_ *‘*s across multiple ***x***_*i*_ *‘*s are more likely to be included in the population-level selection set. For a similar setting, Dai et al. ^49^ showed that under certain additional assumptions, *ℛ*_population_ obtained asymptotically controls the FDR. In our simulations, we observed that the inclusion-rate-based procedure empirically controls the FDR.

#### F. Implementation details of the replicability-improving strategy

To reduce the variation introduced by the randomness of knockoff generation and model fitting, we propose a replicability-improving strategy as described in Section 3.E, following Dai et al.^49^, Specifically, we generate *M* selection sets 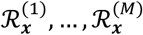 for the target individual with features ***X*** = ***x*** under the FDR level *α* by performing *M* runs of our procedure. The derandomized selection set 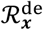 is then computed by applying the inclusion-rate-based procedure as follows:

1. For each run *m* = 1, …, *M*, compute the inclusion rates corresponding to *H*_*j*_ *‘*s from 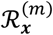:

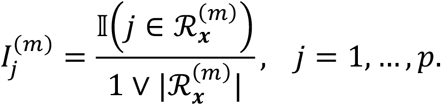
2. Compute the final inclusion rates by averaging across the *M* runs:

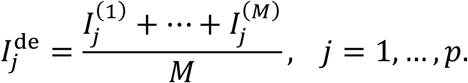
3. Obtain the derandomized selection set 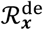 as

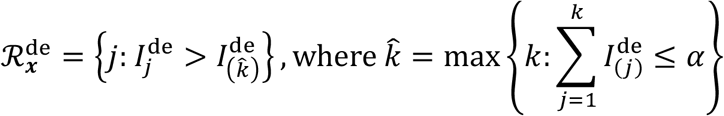

and 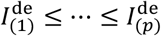 denote the order statistics of 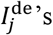.

The derandomized selection set 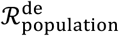 at the population level can also be obtained using an analogous procedure. By averaging the results across multiple runs, the effect of randomness in knockoff generation and model fitting is reduced, resulting in more reproducible and stable selection sets 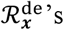 and 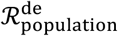. As shown in our simulations, this procedure empirically controls the FDR.

#### G. Additional details for Section 2.F

To quantify the stability and replicability of the relevant procedures, we perform the following experiments:

1. Without derandomization: for *t* = 1, …, 50, perform our localized feature selection procedure to obtain (a) the selection sets 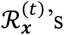 for evaluation points and (b) the population selection set 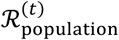 obtained by applying the knockoff filter ^11^ to 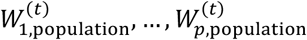 with

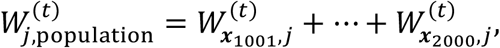

where ***x***_1001_, …, ***x***_2000_ represent 1000 evaluation points.
2. With derandomization: for *t* = 1, …, 50, perform our localized feature selection procedure with the replicability-improving strategy proposed in Section 3.E (with *M* = 15 runs) to obtain the derandomized selection sets 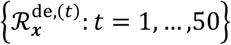 and 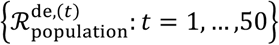.

To quantify the stability of the derandomized localized selection set 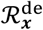 with respect to an evaluation point ***x***, we first compute the average pairwise Jaccard similarity among 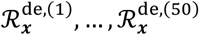 within each training datasets and compute the mean of the 100 average pairwise Jaccard similarities over the 100 training datasets as the stability measure 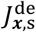. Analogously, we compute the average pairwise Jaccard similarity among 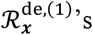 obtained from different training datasets as the replicability measure 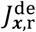. The computational processes of both measures are visualized in Supplementary Figure 2. Corresponding measures with respect to localized selection set *ℛ*_***x***_, population selection set *ℛ*_population_, and derandomized population selection set 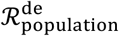 are also computed in an analogous manner.

**Supplementary Figure 2.**
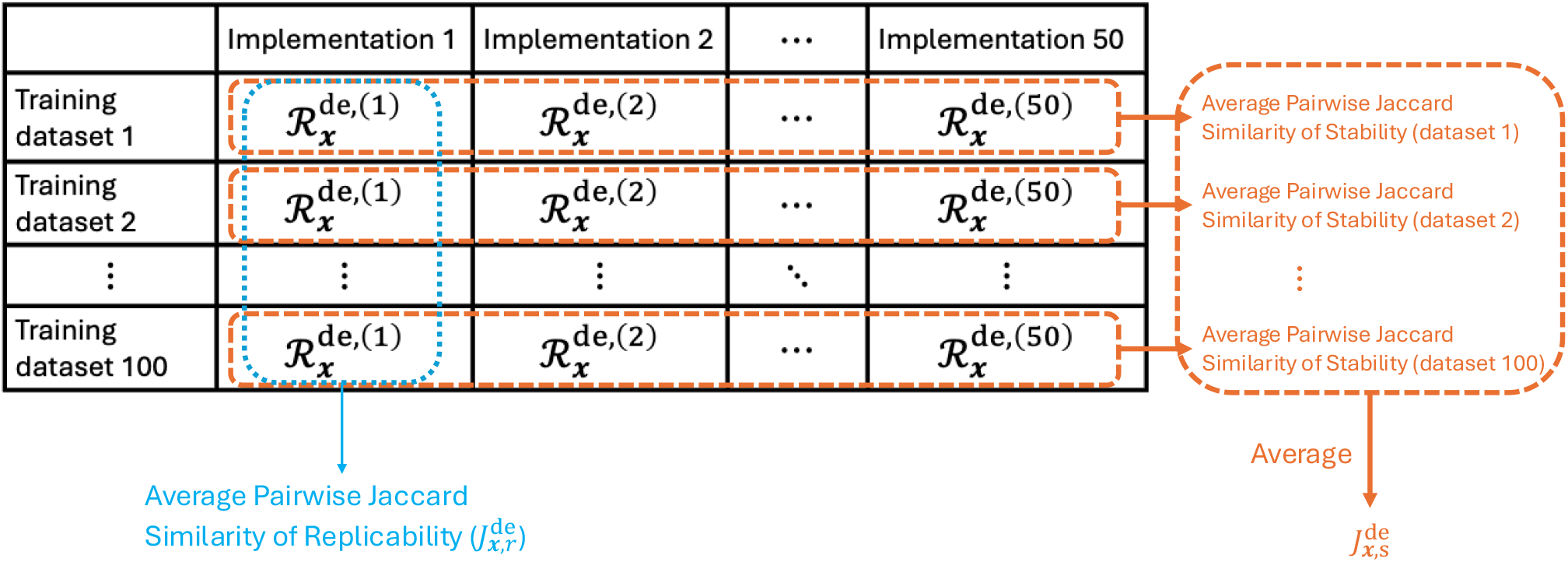
Graphical illustration of the computation processes of the stability measure 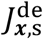 and the replicability measure 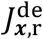.

### S2. Additional simulation studies

#### A. Simulation results for alternative aggregation methods

To evaluate the empirical performance of the three proposed aggregation methods in Section S1.E, we applied them to the simulated datasets generated in Section 2.C. The empirical FDR and power of the population selection set *ℛ*_population_ are computed by averaging the FDP and power over 100 replicates, as shown in Supplementary Figure 3. For comparison, we also include the empirical FDR and power of (a) the population selection set *ℛ*_lasso_, obtained via the model-X knockoff filter using the lasso coefficient-difference statistic, and (b) the union of *ℛ*_***x***_’s (*ℛ*_union_). Specifically, *ℛ*_lasso_ is obtained from a *population feature selection* method with rigorous FDR control, while *ℛ*_union_, without FDR control, serves as an information-based upper bound on power.

**Supplementary Figure 3.**
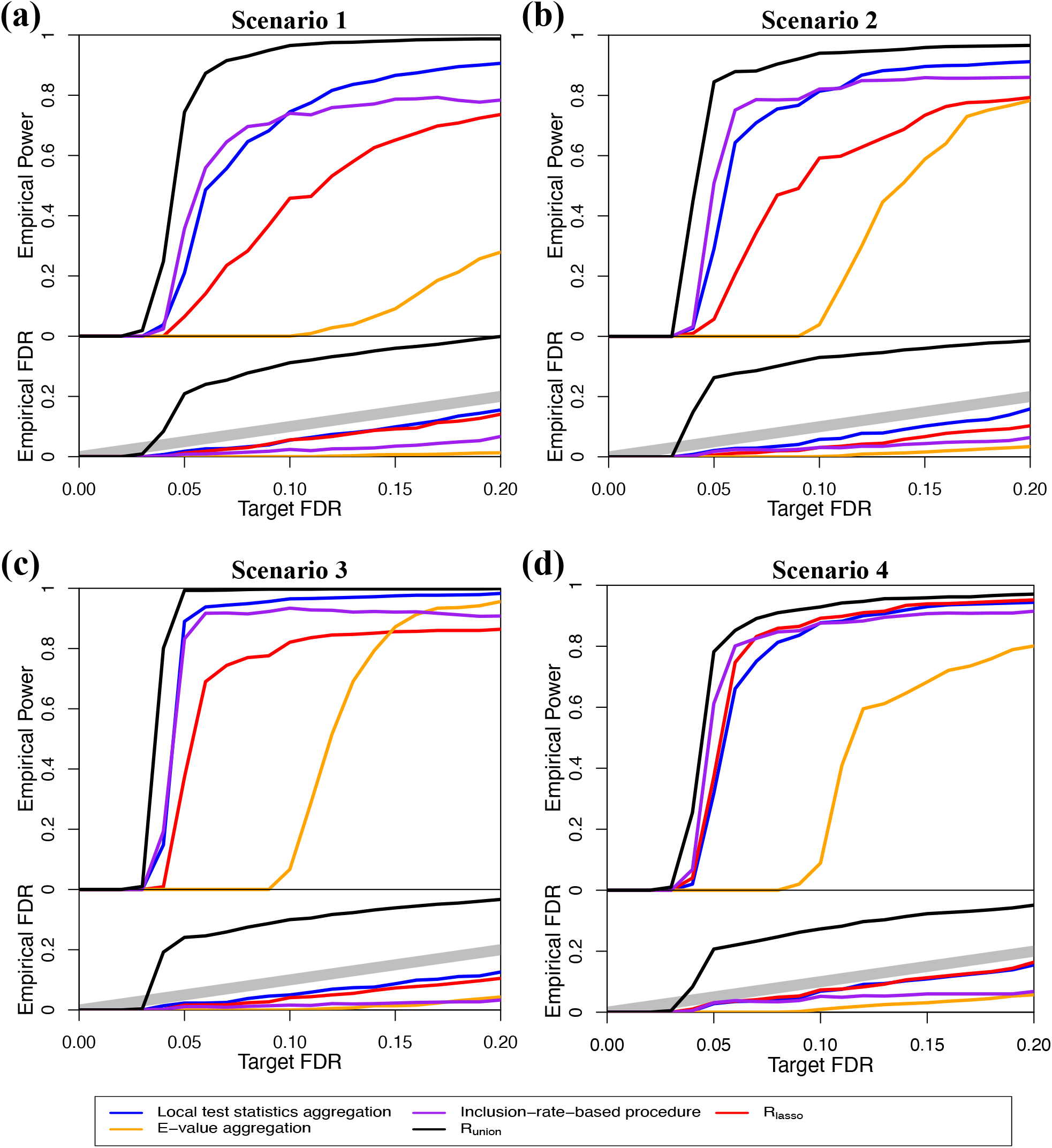
Empirical power and FDR of *ℛ*_population_ using different aggregation approaches under various scenarios with comparison to *ℛ*_lasso_ and *ℛ*_union_.

From Supplementary Figure 3, we observe that all three proposed aggregation approaches effectively control the empirical FDR, regardless of the target FDR level. However, compared to the population selection set *ℛ*_lasso_, the power of these approaches shows varying patterns.

The e-value aggregation technique (Ren & Barber, 2023) exhibits significantly lower power than *ℛ*_lasso_ across all scenarios. In contrast, the population selection sets obtained through either aggregating test statistics *W*_*j*,population_*’*s or the inclusion-rate-based procedure ^49^ demonstrate non-inferior power. Notably, these two approaches generally outperform *ℛ*_lasso_ by a significant margin, except in Scenario 4, where individual heterogeneity is absent. This highlights the advantage of aggregation in capturing heterogeneous localized evidence against false *H*_*j*_’s.

In general, the power of the population selection set obtained by aggregating test statistics *W*_*j*,population_*’*s increases steadily with higher target FDR levels. On the other hand, the power of the inclusion-rate-based procedure remains unchanged when the target FDR level exceeds 0.1. This behavior can be attributed to the non-nesting property of the inclusion-rate-based procedure across different target FDR levels, potentially limiting its asymptotic power. In contrast, the population selection set produced by the aggregated test statistics *W*_*j*,population_*’*s does not suffer from this issue. Additionally, the power curve for aggregating localized test statistics *W*_***x***,*j*_’s is closer to the power curve of *ℛ*_union_ (the information-based upper bound of power). This is because localized test statistics *W*_***x***,*j*_ *‘*s are more informative in capturing the strength of evidence against *H*_*j*_ *‘*s, suggesting that aggregating localized test statistics *W*_***x***,*j*_’s is a more effective approach.

#### B. Simulation results for a neural network model

In this section, we present the results when a neural network model *h*(⋅) is fitted on the extended dataset 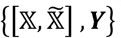 under Scenario 3 in Section 2.C. In our implementation, the neural network model *h*(⋅) contains a single fully connected hidden layer, whose dimension is selected from {2,5,8,10,15,20,30,50,103} based on a 20% validation dataset. Following the hidden layer, we include a batch normalization step, a ReLU activation function, a dropout (rate = 0.5) and a linear output layer. In addition, lasso penalty is applied on network parameters, where the penalty strength is also selected based on the same 20% validation dataset as above. The training will be terminated when the validation loss does not improve for 10 consecutive epochs.

Supplementary Figure 4 (a) shows the average power and FDR of *ℛ*_***x***_’s, across 100 replicates with neural network model *h*(⋅). Compared to Figure 3 (c), the neural network model is less effective in capturing localized heterogeneity. Although the effect size of important features grows with respect to age across all subgroups, the power of *ℛ*_***x***_ *‘*s for AFA females does not increase significantly. This is further illustrated in Supplementary Figure 4 (b), where the UMAP projections for different subgroups do not exhibit clear separation, unlike Figure 4 (c). This is mainly due to the small sample size that causes the neural network model to overfit and capture more of the population-level effect rather than finer localized effects for different subgroups.

**Supplementary Figure 4.**
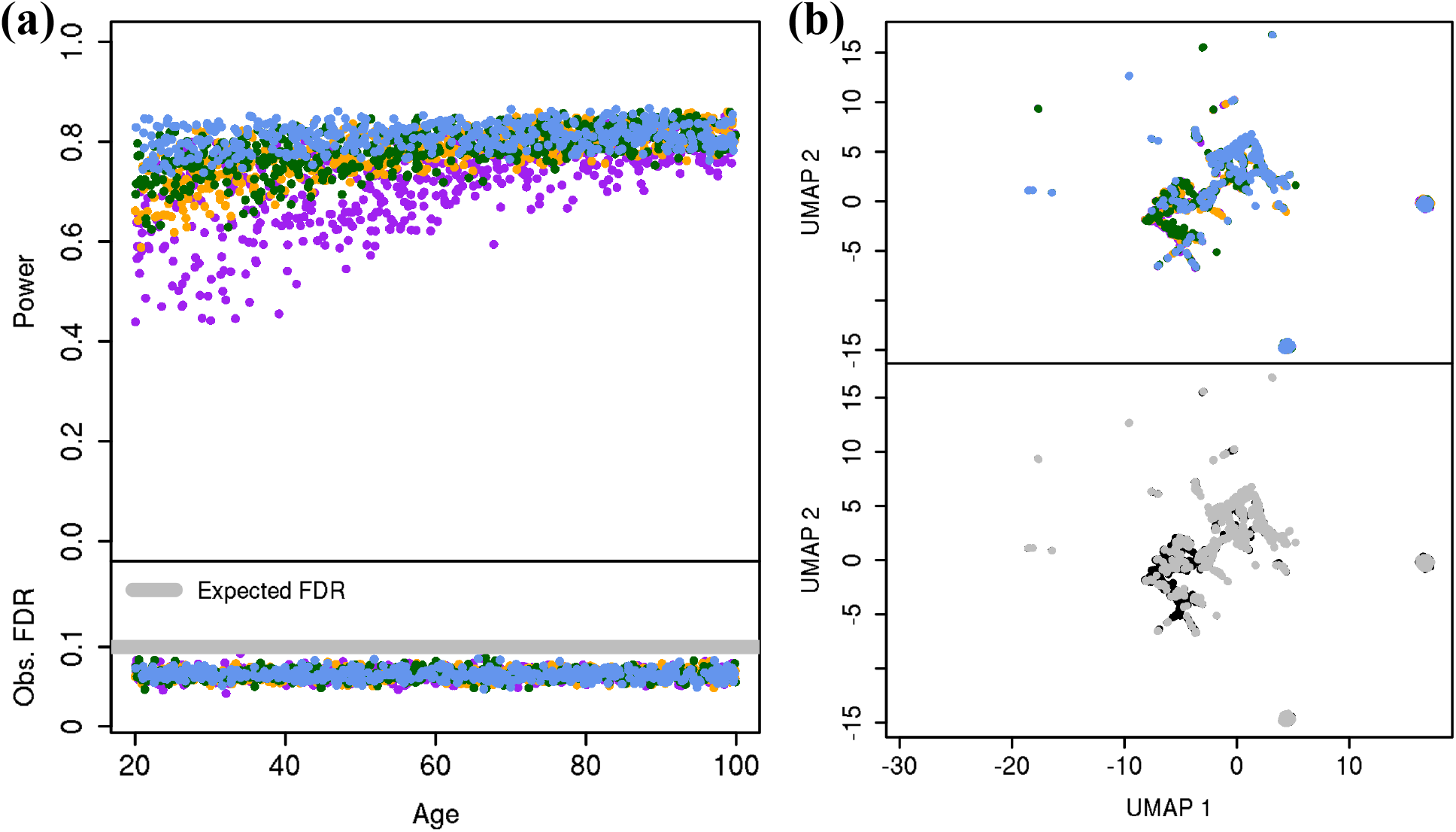
(a) Average power and FDP across 100 replicates of the localized selection sets *ℛ*_***x***_’s (with respect to the target FDR level *α* = 0.1) for each of the 2000 evaluation points under Scenario 3. (b) UMAP visualization of the augmented selection vectors ***v***_***x***_’s from one replicate. The upper panel of each subfigure is colored according to ancestry and gender profiles (*E*_1_, *E*_2_), while the lower panel is colored according to whether the age exceeds 60 (the median of age).

### S3. Additional results for ACRC scRNA-seq data analysis

**Supplementary Table 1.** Number of microglia in each individual and brain region.

| Individual | Cortex | Hippocampus | Individual | Cortex | Hippocampus |
| --- | --- | --- | --- | --- | --- |
| AD case 1 | 0 | 166 | Control 1 | 64 | 117 |
| AD case 2 | 0 | 127 | Control 2 | 0 | 64 |
| AD case 3 | 0 | 181 | Control 3 | 0 | 53 |
| AD case 4 | 160 | 160 | Control 4 | 0 | 95 |
| AD case 5 | 109 | 87 | Control 5 | 0 | 93 |
| AD case 6 | 176 | 55 | Control 6 | 69 | 67 |
| AD case 7 | 77 | 84 | Control 7 | 98 | 89 |
| AD case 8 | 0 | 17 | Control 8 | 94 | 239 |
| AD case 9 | 0 | 832 |  |  |  |

**Supplementary Figure 5.**
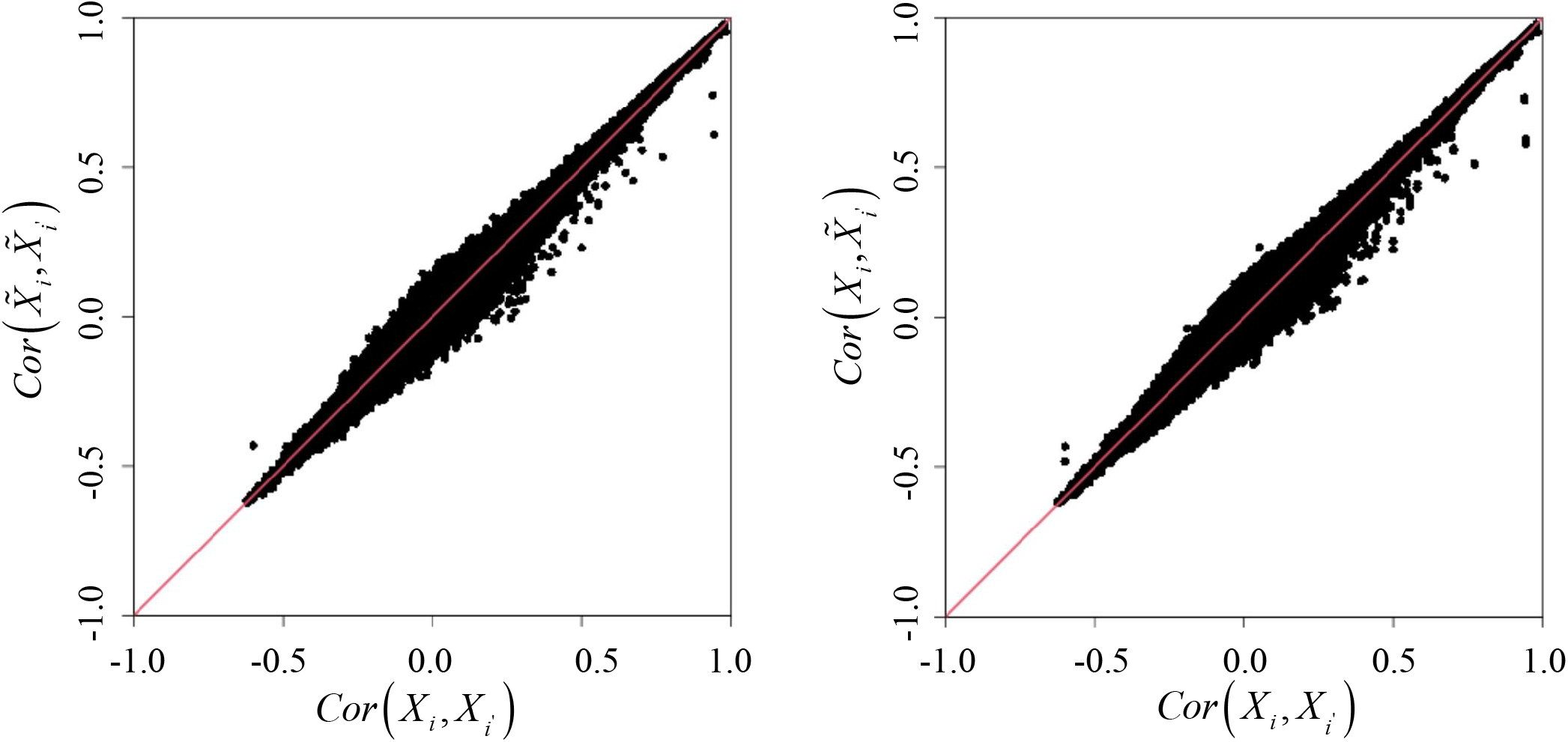
Comparisons of empirical correlations between *X*_*j*_ and *X*_*k*_ with the empirical correlations between 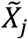 and 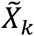 (left panel), and comparisons of correlations between *X*_*j*_ and *X*_*k*_ with the empirical correlations between *X*_*j*_ and 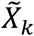(right panel) under the ACRC scRNA-seq data.

**Supplementary Figure 6.**
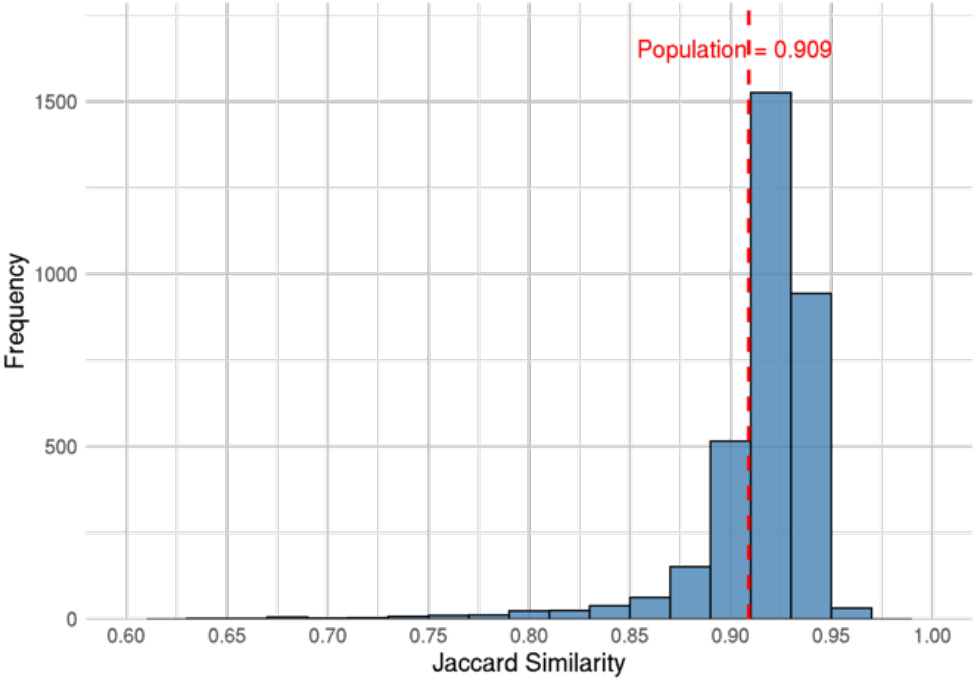
Histogram of the average pairwise Jaccard similarities over 30 runs for the 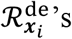. The red dashed line indicates the corresponding stability measure for 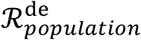. Both are computed under the ACRC scRNA-seq data.

**Supplementary Table 2.** Genes included in 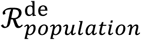 and their corresponding inclusion rates under the ACRC scRNA-seq data.

| Gene | Inclusion Rate | Gene | Inclusion Rate | Gene | Inclusion Rate |
| --- | --- | --- | --- | --- | --- |
| FOXP1 | 4.025% | JAZF1 | 2.906% | ADAMTS9 | 2.453% |
| MAP3K8 | 3.775% | HDAC9 | 2.859% | FLT1 | 2.329% |
| MALAT1 | 3.568% | MRC1 | 2.841% | SLC2A3 | 2.180% |
| DICER1 | 3.507% | ZCCHC2 | 2.808% | VTI1A | 2.067% |
| HSPB1 | 3.434% | FP236383 | 2.759% | EGFL7 | 1.941% |
| BSG | 3.315% | PRKCA | 2.722% | HIVEP3 | 1.747% |
| SLC7A5 | 3.250% | PCDH9 | 2.636% | ACSL1 | 1.734% |
| ETV6 | 3.221% | FP671120 | 2.552% | NEAT1 | 1.711% |
| SLC22A23 | 3.132% | PKM | 2.511% | SLC11A1 | 1.641% |
| XIST | 3.068% |  |  |  |  |

